# “Upregulation of TLR4/MyD88 pathway in alcohol-induced Wernicke’s encephalopathy: findings in preclinical models and in a postmortem human case”

**DOI:** 10.1101/2022.06.30.497714

**Authors:** Marta Moya, Berta Escudero, Elena Gómez-Blázquez, Ana Belen Rebolledo-Poves, Meritxell López-Gallardo, Carmen Guerrero, Eva María Marco, Laura Orio

## Abstract

Wernicke’s encephalopathy (WE) is a neurologic disease caused by vitamin B1 or thiamine deficiency (TD), being the alcohol use disorder (AUD) its main risk factor. WE patients present limiting motor, cognitive and emotional alterations related to a selective cerebral vulnerability. Neuroinflammation has been proposed as one of the phenomena contributing to brain damage. Our previous studies provide evidence for the involvement of the innate immune receptor Toll-like (TLR) 4 in the inflammatory response induced in the frontal cortex and cerebellum in TD animal models (animals fed with TD diet and receiving pyrithiamine). However, the effects of the combination of chronic alcohol consumption and TD on TLR4 and their specific contribution to the pathogenesis of WE are currently unknown. Additionally, no studies on TLR4 have been conducted on WE patients since brains from these patients are difficult to achieve. Here, we used rat models of chronic alcohol (CA; 9 months of forced consumption of 20% (w/v) alcohol), TD hit (TDD; TD diet + daily 0.25 mg/kg i.p. pyrithiamine during 12 days), or a combined treatment (CA+TDD) to check the activation of the proinflammatory TLR4/MyD88 pathway and related markers in the frontal cortex and the cerebellum. In addition, we characterized for the first time the TLR4 and its co-receptor MyD88 signature, along with other markers of this proinflammatory signaling such as phospo-NFκB p65 and IκBα, in the post-mortem human frontal cortex and cerebellum (gray and white matter) of an alcohol-induced WE patient, comparing it with negative (no disease) and positive (aged brain with Alzheimer’s disease) control subjects for neuroinflammation. We found an increase in the cortical TLR4 and its adaptor molecule MyD88, together with an upregulation of the proinflammatory signaling molecules p-NF-ĸB and IĸBα in the CA+TDD animal model. In the patient diagnosed with alcohol-induced WE we observed cortical and cerebellar upregulation of the TLR4/MyD88 pathway. Thus, our findings provide evidence, both in the animal model and the human postmortem brain, of the upregulation of the TLR4/MyD88 proinflammatory pathway in WE related to alcohol consumption.

## INTRODUCTION

Wernicke’s encephalopathy (WE) and Korsakoff’s syndrome are considered different stages of the same disease due to vitamin B1 or thiamine deficiency (TD), where WE represents the acute and reversible (when treated with thiamine) form of the disease and Korsakoff is an advanced and irreversible state characterized by neuronal death. This neurologic disease, also named Wernicke-Korsakoff syndrome (WKS), is characterized by ocular abnormalities (nystagmus and/or ophthalmoplegia), mental status changes, and gait disturbances (Kohnke and Meek, 2021). Because of limiting motor, cognitive and emotional alterations, these patients require heavy dependence to complete daily life activities.

Alcohol use disorder (AUD) is the main risk factor for this disease, although other causes with no history of alcohol dependence may also induce the pathology, as repetitive vomiting, gastric disorder, or after bariatric surgery (Kopelman, 1995; Deb et al., 2001). The nutritional TD in AUD is associated with malnourishment and decreased absorption of thiamine, due to direct effects of alcohol on its metabolism, besides reduced storage in the liver because of an alcoholic liver disease (Arts et al., 2017).

The bioactive form of thiamine (thiamine diphosphate) is necessary for energy metabolism in all cells. Therefore, the brain is the main site of TD-induced damage due to its immense energy requirement compared to the rest of the body (Clarke and Sokoloff, 1999). Brain damage has been extensively described in several brain regions in WE, mainly including diencephalic regions such as the thalamus and mammillary bodies (Manzo et al., 2014). However, some authors pointed out to the presence of damage in other structures less studied in this pathology, such as the frontal cortex (Jacobson and Lishman, 1990; Jernigan et al., 1991; Paller et al., 1997; Aupée et al., 2001; Gibson et al., 2016) and the cerebellum (Mulholland, 2006; Manzo et al., 2014). In our previous preclinical studies about WE, we selected these less studied frontal cortex and cerebellum (Moya et al., 2021, 2022) as structures of great interest to be investigated, since both participate in motor function control, in cognition and in emotional responses (Baillieux et al., 2008; Molinari et al., 2008; Rudebeck et al., 2008; Leggio et al., 2011; Clausi et al., 2017). Indeed, the frontal cortex is particularly important in executive control tasks, behavioral inhibition, including cognitive processes, social behavior and inhibition of motor responses. Within the cerebellum, the mapping of associative learning with emotional, motor, and cognitive functions follows a medial-to-lateral cerebellar distribution: the sensorimotor functions are distributed more toward the midline, while the cognitive functions are located more laterally in the cerebellar hemispheres. Executive functions, including verbal working memory, are related to both cerebellar hemispheres, while affective functions are primarily midline in the so called “limbic cerebellum”. Interestingly, the left cerebellar hemisphere, region analyzed in the present study, also appears to be involved in visuospatial functions and in linguistic processes (Klein et al., 2016; Amore et al., 2021). Thus, the cerebellum and the frontal cortex are two brain areas directly involved in the behavioral alterations manifested in the WE, which deserve further investigation.

The exact cause of brain damage in WE is unclear, but neuroinflammation has been proposed as a contributing factor (Neri et al., 2011) (Zahr et al., 2014; Toledo Nunes et al., 2019). Proinflammatory cytokines, enzymes and different constituents of this process have been reported, but how the inflammatory response is activated in the brain tissue remains unknown. Very recently, our research group reported for the first time the involvement of the innate immune receptor Toll-like (TLR)-4 in the pathogenesis of non-alcoholic WE, showing a selective vulnerability of the frontal cortex and cerebellum, two brain structures understudied in comparison to diencephalic regions, in this pathology over time (Moya et al., 2021).

The activation of the canonical proinflammatory TLR4 pathway induces, *via* Myeloid Differentiation factor 88 (MyD88), the recruitment of downstream signaling molecules that triggers the stimulation of transcriptional factors, such as the nuclear factor κB (NF-κB), which lead to the induction of genes encoding inflammation-associated molecules and cytokines. In addition to cytokines, NF-κB transcriptional activity induces the expression of other pro-inflammatory markers that lead to oxidative and nitrosative stress, such as the inducible nitric oxide synthase (iNOS) and cyclooxygenase 2 (COX-2) enzymes, and different caspases, generating lipid peroxidation and apoptotic cell death, respectively. Some other molecules can be released in response to injured tissue, such as heat shock proteins (HSPs) and the high mobility group box 1 protein (HMGB1), inducing more neuroinflammation in a vicious cycle [reviewed in (Orio et al., 2019)]. The TLR4-induced neuroinflammatory pathway has been extensively studied in the context of AUD (Pascual et al., 2011; Crews et al., 2013; Montesinos et al., 2016; Antón et al., 2017) and we recently reported that the TLR4-induced neuroinflammation in the frontal cortex and cerebellum in TD animals could be related to the cognitive and motor deficits, respectively (Moya et al., 2021). However, the specific contribution of TD and chronic alcohol use in the impact of TLR4 signaling and their contribution to the pathogenesis of WE are currently unknown.

In the present study we aimed to further characterize the role of the TLR4 in WE, by using combined models with TD and chronic alcohol exposure, the two main known contributing factors of the pathology, and we also explored the TLR4 activation and signalling in the frontal cortex and cerebellum of a postmortem alcohol-induced WE brain. The presence of postmortem brains of WE-diagnosed patients in biobanks is extremely scarce. Here, we reported a deep analysis (in white and grey matter in frontal cortex and cerebellum) in a single case, using a matched control subject and a positive control in which TLR4-induced neuroinflammation has been extensively reported, as in an aged brain with Alzheimer’s disease.

## MATERIALS AND METHODS

### Rodent Studies

#### Animals and housing

Male Wistar rats (Envigo©, Barcelona, Spain) (n = 50), weighing 100-125 g at arrival were used. Animals were housed in groups of 2–3 per cage and maintained at a constant room temperature (21 ± 1^°^C) and humidity (60 ± 10%) in a reversed 12 h dark-light cycle (lights on at 8:00 p.m.). Standard food and tap water were available *ad libitum* during an acclimation period of 12 days prior to experimentation and then rats were randomly assigned to the experimental groups.

All procedures followed ARRIVE guidelines and were adhered to the guidelines of the Animal Welfare Committee of the Complutense University of Madrid (reference: PROEX 312-19) in compliance with the Spanish Royal Decree 118/2021 and following the European Directive 2010/63/EU on the protection of animals used for research and other scientific purposes.

#### Experimental Groups

The experimental design and all the procedures of this animal study are described in detail and can be viewed at (Moya et al., 2022).

Briefly, to explore the different conditions that contribute to develop WE, the following experimental groups were used:

##### Chronic alcohol (CA)

animals exposed to forced consumption of 20% (w/v) alcohol for 9 months (*n* = 9).

##### TD diet (TDD)

TD hit (TD diet* + pyrithiamine 0.25 mg/kg dissolved in saline (0.9% NaCl) i.p. daily injections the last 12 days of experimentation; *TD Diet specific composition is detailed in the Supplementary Material) (*n* = 9).

##### Chronic alcohol combined with TD diet in the last days of treatment (CA + TDD)

both combined treatments (*n =* 10).

These groups were compared with the corresponding *control group (C)*, animals drinking water with standard chow (*n* = 8).

During the last 12 days of TDD protocol, the remaining animals (C and CA) received equivalent daily injections of vehicle (saline, i.p.).

The number of animals in the alcohol and TDD groups was slightly higher to control groups for the possible loss of experimental subjects.

We consider that the group with the combined CA+TDD treatment is the most relevant in this translational study, since is the animal model that most closely approximates the WE related to alcohol use.

#### Tissue samples collection

The day 12 of TDD protocol, at least 1 h after treatment administration, all animals were killed by rapid decapitation after anesthesia overdose of sodium pentobarbital (320 mg/kg, i.p., Dolethal®, Vétoquinol, Spain). Brains were immediately isolated from the skull, discarding meninges and blood vessels, and the frontal cortex (area between Bregma +4.7 and + 1.2 mm aprox.) and the left cerebellar hemisphere were dissected on ice and frozen at - 80°C until assayed. The liver was also immediately taken out and kept at -80°C for other assays.

#### Western Blot Analysis

Frontal cortex and cerebellar hemisphere samples were processed and analyzed by western blot following the methodology previously detailed in (Moya et al., 2022).

Briefly, the tissue samples were homogenized at a ratio of 1:3 (w/v) in ice-cold lysis-buffer with protease inhibitors, followed by centrifugation to obtain the supernatants. Protein levels were measured by Bradford’s method (Bradford, 1976). The samples were adjusted with the loading buffer to a final concentration of 1 mg/mL, and 15–20 µg of total protein were separated by SDS-polyacrylamide gels and transferred to nitrocellulose membranes. Blots were incubated with specific primary and secondary antibodies, using the housekeeping ß-actin protein as loading control (see Table 1 for complete list of antibodies and their details). Bands were visualized using an ECL kit and quantified by densitometry using ImageJ software (NIH, USA).

**TABLE 1.** Specific antibodies used in Western blotting to detect proteins of interest.

| *Protein* | *Primary antibody<sup>1</sup>* |  | *Secondary antibody* |
| --- | --- | --- | --- |
| TLR4 | *sc-293072* | 1:500 BSA 1% | Mouse (1:2000) |
| MyD88 | *ab2064* | 1:750 BSA 5% | Rabbit (1:2000) |
| phospho-NFκB p65 | (27.Ser 536) *sc-136548* | 1:400 | Mouse (1:2000) |
| NFκB p65 | *sc-372* | 1:1000 | Rabbit (1:2000) |
| IκBα | *sc-371* | 1:1000 BSA 5% | Rabbit (1:2000) |
| COX-2 | *sc-376861* | 1:750 BSA 2% | Mouse (1:2000) |
| iNOS | *sc-650* | 1:500 BSA 2% | Rabbit (1:2000) |
| HSP70 | *sc-1060* | 1:1000 | Goat (1:4000) |
| β-actin | *A5441 Sigma* | 1:10000 | Mouse (1:10000) |

<sup>1</sup> References (codes) and dilutions. Abbreviations: [TLR4: Toll-like receptor 4; MyD88: myeloid differentiation factor 88; Phosphorylated p65 subunit of NF κB: nuclear factor kappa B; IκB $\alpha$ : I kappa B alpha protein; COX-2: cyclooxygenase-2; iNOS: inducible oxide nitric synthase; HSP70: heat shock protein 70; BSA: Bovine serum albumin; sc: Santa Cruz Biotechnology; ab: Abcam]. Sources of secondary antibodies: Anti-Mouse IgG, HRP-linked whole Ab (from sheep) (GE Healthcare, ref. NXA931); Anti-Goat IgG (whole molecule)–Peroxidase antibody produced in rabbit (Sigma-Aldrich, ref. A5420); Anti-Rabbit IgG (H+L) Cross-Adsorbed Secondary Antibody, HRP conjugate (from donkey) (ThermoFisher Scientific, ref. 31458).

#### Liver damage

The status of the liver in the animals was checked by measuring the hepatic nitrites and malondialdehyde (MDA) levels, due to the major role of these processes in the pathogenesis of alcohol liver disease (ALD) (McKim et al., 2003; Galicia-Moreno and Gutiérrez-Reyes, 2014; Pérez-Hernández et al., 2017; Tan et al., 2020; Yang et al., 2022). For details, see Suppementary Data 1.5.

### Postmortem Human Studies

#### Cases

Three cases were selected from brains donated to the Biobank of the Hospital Universitario Fundación Alcorcón (HUFA), Madrid, Spain:

##### Case diagnosed with WE

woman, 62 years old. History of chronic alcohol consumption of at least 1 bottle of wine per day for 10-15 years. The patient showed classical symptoms of WE: altered mental state such as confusional syndrome, disorientation in space and time, scarce and incoherent spontaneous language; ocular signs (horizontal nystagmus) and motor disturbances (extrapyramidal symptoms, decreased reflexes). The liver enzymes’ (transaminases) values and other information is reported in Supplemental Data, section 1.1. The WE was diagnosed on the basis of the (previous) clinical presentation along with the confirmation by postmortem neuropathological analyses.

##### *Negative control cas*e

woman, 53 years old. Cause of death due to a non-neurological disease or psychiatric disorders.

##### Positive control case: aged brain with Alzheimer’s disease (AD)

woman, 76 years old. Primary progressive aphasia, logopenic subtype. Frontotemporal lobar degeneration. Neuropathological diagnosis with changes of AD advanced stage. This was the positive control to observe neuroinflammation, since it had a double hit: aged brain with Alzheimer’s disease. (It has already been demonstrated that TLR4-neuroinflammation is involved in AD pathology (Zhang et al., 2012; Fiebich et al., 2018; Miron et al., 2018; Calvo-Rodriguez et al., 2020) and an aged brain is also susceptible of having more neuroinflammation).

The extended clinical history of each case can be found in the Supplementary Materials and Methods, section 1.1.

#### Sample processing

Postmortem proceedings were carried out in the Hospital Universitario Fundación Alcorcón (HUFA), Madrid, Spain. All studies were performed complying with national ethical and legal regulations, being approved by the Drug Research Ethics Committee of the HUFA (ref 62-2018).

According to the brain bank protocol, in conventional donation cases, immediately after extraction, the left half of the brain was fixed by immersion in phosphate-buffered 4% formaldehyde for at least 3 weeks. Then, the brain is processed by coronal slices, except for the cerebellum, which is sectioned sagittally. Brain samples from the dorsolateral frontal cortex (Brodmann area 9) and the left cerebellar hemisphere (corresponding to the area from the superior cerebellum to the dentate nucleus) were selected for this study. The tissue was embedded in paraffin and 4μm sections were obtained by microtomy for subsequent immunostaining.

#### Immunohistochemistry (IHC)

The detailed description of the IHC protocol is provided in the Supplementary Materials and Methods, section 1.2. Briefly, slides were incubated with specific primary antibodies against TLR4, MyD88, p-NFκB p65 and IκB-α, and were developed by diaminobenzidine (DAB) along with Carazzi’s hematoxylin as counterstaining.

To evaluate the specificity of the staining, several technical controls were run including, on one hand, the omission of primary antibody and, on the other hand, omission of the secondary antibody. These technical controls resulted in the absence of staining, and they were performed in both frontal cortex and cerebellar tissue from the control, alcohol-induced WE and AD cases. In addition, the specificity of the TLR4 and MyD88 antibodies selected for this study was previously demonstrated in human brain tissue by using IHC and western blotting (Zurolo et al., 2011; MacDowell et al., 2017; Martín-Hernández et al., 2018).

#### Imaging and Quantification

Slides immunostaining of the frontal cortex and cerebellar hemisphere were observed under light microscopy (Zeiss Axioplan Microscope, Germany). The microscope had a high-resolution camera attached (Zeiss Axioplan 712 color, Germany), which was used for capturing the images which were then processed using Axiovision 40V 4.1 (Carl Zeiss vision, Germany) and ZEN2 software (Carl Zeiss AG, Oberkochen, Germany). Light, shine and contrast conditions were kept constant during the capture process. For the study of each tissue section per patient, a total of 16 visual fields, 8 within the gray matter and another 8 within the white matter, were examined. An image of each visual field was taken at 40x magnification for the frontal cortex, and at 20x for the cerebellum in order to capture its three layers.

In addition, a manual neuronal counting of each image was performed, obtaining a total number range for an accurate comparison between the cases.

Positive signal (in brown color due to DAB) on immunohistochemically stained tissues were semi-quantitatively evaluated by a visual and an automatic scoring, comparing both methods to achieve the most reliable results. Images were always evaluated in a blinded manner without prior knowledge of the clinical information.

##### Visual/observational analysis

immunopositivity of the images was visually assessed by the investigator using a scoring system adapted to our study. The modified immunoreactivity score (IRS) is a composite score assigned to distribution and intensity of immunostaining, based on (Wang et al., 2011) and (Meyerholz and Beck, 2018) (see Supplementary Data, section 2.1, Figure 1). Briefly, the observer must assign sub-scores for immunoreactive distribution (on a 0–4 scale) and intensity (on a 0–3 scale), multiplying them to calculate the total score for each image (ranged from 0 to 12). Final IRS was obtained by averaging the values in the eight fields for each section.

**FIGURE 1.**
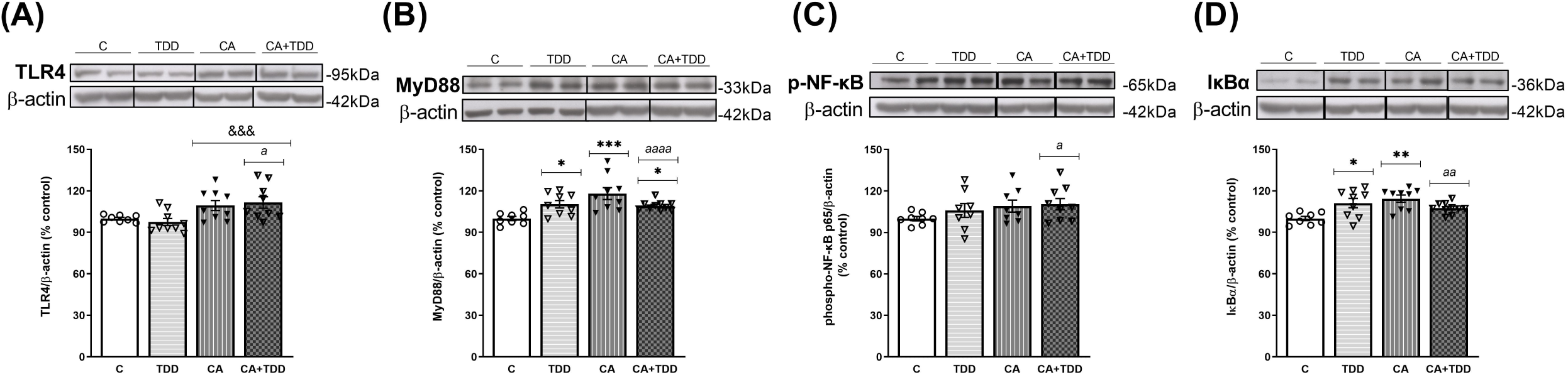
Effects in the TLR4 signaling pathway in Frontal Cortex of TDD, CA and CA+TDD-treated rats. Graphs indicate protein levels of (A) TLR4, (B) MyD88, (C) p-NF-ĸB and (D) IĸBα markers by Western blot; data of the respective bands of interest (upper bands) were normalized by β-actin (lower band) and expressed as a percentage of change versus the control group. Some blots were cropped from the original (black lines) for improving the clarity and conciseness of the presentation. Mean ± SEM (n = 8-10). Two-way ANOVA or nonparametric Kruskal-Wallis test. Overall alcohol effect: &&&p < 0.001; different from control group: *p < 0.05, **p < 0.01, ***p < 0.001. Since the combined CA+TDD treatment mimics better the human case of alcohol-induced WE, this group was also compared with the C group by Unpaired Student’s t-test or Mann-Whitney (CA+TDD vs C): a p < 0.05, aa p < 0.01, aaaa p < 0.0001.

##### Automated analysis

a semiquantitative analysis of images was carried out using the *ImageJ Fiji* software, following the color deconvolution protocol previously described by (Crowe and Yue, 2019). In brief, a threshold value was set to remove the background signal after the deconvolution of images, followed by the quantification of the DAB signal within the image. The average intensity of DAB signal in IHC images was calculated. Finally, the mean value of the eight images was taken to represent the specific immunoreactivity of each target protein.

Since valuable information might be neglected by the above-mentioned scoring systems, we included a brief description of the cell types and tissue components positively marked, as well as about the intensity and characteristics of the staining (see Supplementary Materials and Methods, section 1.3).

Both visual IRS and automatic *Fiji* methods were employed for the analysis of frontal cortex images. Since both procedures reported comparable and reliable results (see results section), the cerebellar hemisphere was subsequently analyzed only by the *Fiji* method.

### Statistical Analysis

Data are expressed as mean ± S.E.M. In the animal study, two-way ANOVAs were used to assess the overall effects or interactions between two factors: CA and TDD. Additionally, unpaired Student’s t-test was used to compare the CA+TDD group *vs* control. Regarding immunohistochemistry analyses, automated *Fiji* measures were analyzed by one-way ANOVA. For manual IRS data the non-parametric Kruskal–Wallis test followed by paired comparisons with Mann-Whitney test was used. Comparison between manual and automated IHC measurements was performed by Pearson’s correlation and linear regression analyses. Parametric tests were performed when normality and homoscedasticity were verified (checked by Kolmogorov-Smirnov and Barlett’s tests, respectively). Otherwise, data were transformed, or the alternative non-parametric analysis was applied. In the ANOVAs, Bonferroni *post hoc* test was used when appropriate. Outliers were analyzed using the Grubbs’ test. A p value <0.05 was set as the threshold for statistical significance in all statistical analyses. The data were analyzed using GraphPad Prism version 8.0 (GraphPad Software, Inc., La Jolla, CA, USA).

## RESULTS

### Frontal cortex findings

#### CA+TDD-treated rats showed an increased expression of TLR4, MyD88, p-NF-ĸB and IĸBα proteins in the frontal cortex

Chronic alcohol increased TLR4 expression levels (Figure 1A, overall effect F _(1, 31)_ =13.7, p=0.0008). Regarding its co-receptor, rats exposed to TDD showed a significant increase in the MyD88 protein levels compared to controls (p=0.0434), being higher in the CA group (p=0.0007 compared to C). Likewise, the combined CA+TDD treatment also induced MyD88 upregulation respect to control animals (p=0.0315) (Figure 1B, differences between groups H=15.82, p=0.0012).

CA exposure induced an increasing trend in the phosphorylation of NF-ĸB that did not reach significance by the ANOVA (Figure 1C, overall effect F _(1, 29) =_2.995, p=0.0941). We report the results of the phosphorylated-NFkB protein normalized by the structural protein β-actin, as done with the rest of the markers, in accordance to other authors (Yang et al., 2021), since the increase of the phosphorylation in p65 subunit is indicative of NF-κB activation to mediate inflammatory gene transcription. The levels of total NFkB were measured, with no changes (see Supplementary Material, section 2.3, Figure 5). We analyzed also the IĸBα protein as a reporter of NF-ĸB activity, finding an increase in its levels by effect of TDD (p=0.042) and CA (p=0.0039) treatments relative to controls (Figure 1D, differences between groups H=12.75, p=0.0052). The increased expression of the NF-κB inhibitory protein IκBα can be considered an autoregulatory mechanism switched on by NF-κB to block its stimulation.

Moreover, cyclooxygenase-2 (COX-2) enzyme was also studied in the frontal cortex, showing an interaction between CA and TDD factors (F _(1, 25)_ =7.407, p=0.0117). *Post hoc* analysis revealed no statistical differences among groups (Supplementary Data, section 2.3, Figure 4A).

In addition, trying to achieve the best approximation between the animal model and the human case, we consider that the combined CA+TDD treatment is an animal model that mimics better the WE related to alcohol use. According to this, we analyzed separately the CA+TDD group by Student’s t-test or Mann Whitney test. CA+TDD group showed higher protein levels of TLR4 and MyD88 compared with controls (Figure 1A, B, U=13, p=0.0274; t=5.208, df=16, p<0.0001, respectively). Additionally, an elevation in p-NF-ĸB and IĸBα protein expression was also observed in this group respect to control (Figure 1C, D, t=2.260, df=15, p=0.0391; t=3.862, df=16, p=0.0014, respectively). No significant changes were observed in COX-2 levels of CA+TDD animals versus control (U=22, p>0.05, n.s.) and other markers (see Supplementary Data, section 2.3, Figure 4).

### Postmortem human frontal cortex of alcohol-induced WE showed an increased expression of TLR4, its co-receptor MyD88 and phospo-NFκB p65

Prior to visual and automatic analysis of the images, the results of the manual counting of the total number of neurons (mainly pyramidal) showed that the three cases studied were within the same range, so they were comparable (data not shown).

The findings reported below were obtained by the automatic *ImageJ Fiji* software, which were confirmed by comparison with the manual IRS analyses. Correlations between both measurements were high (Supplementary Data, section 2.2, Figure 3 and Table 1, for TLR4: r=0.6375; for MyD88: r=0.7958; for both p<0.0001) supporting that the *Fiji* protocol here used is a robust automated measure for TLR4 and MyD88 IHC staining in the brain tissue. In addition, *Fiji* data were chosen as representative results since this method is the most objective. Manual IRS results for frontal cortex images can be found in the Supplementary Data, section 2.2, Figure 2).

**FIGURE 2.**
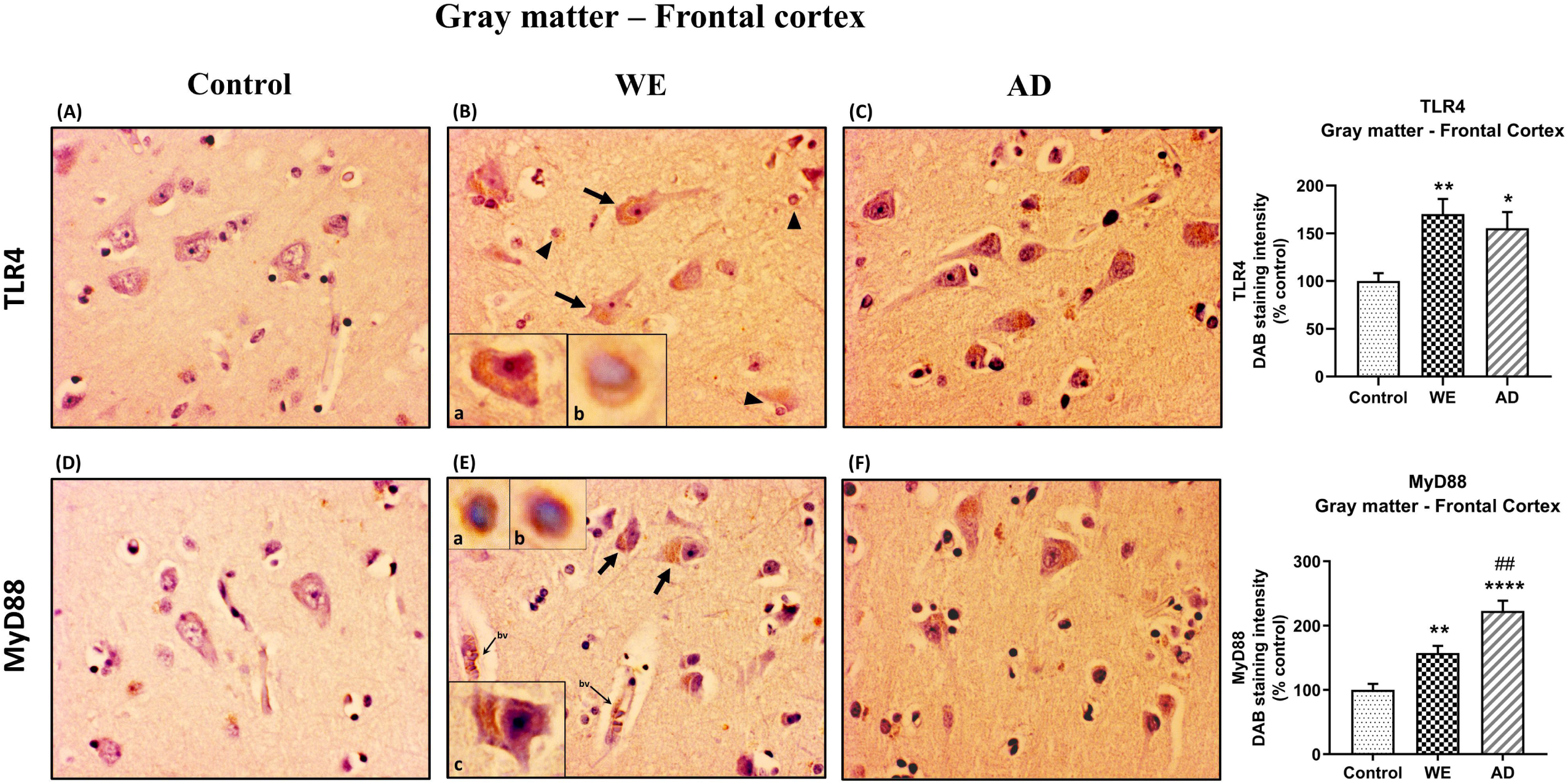
Representative images of immunohistochemical detection for TLR4 and MyD88 in the gray matter of the frontal cortex of control, alcohol-related Wernicke’s encephalopathy (WE) and Alzheimer’s disease (AD) cases. Regarding to TLR4 and MyD88 results in the WE case (B), pyramidal neurons were strongly stained especially in the cytoplasm (arrows; high magnification in inset B-a; E-c) and also glial immunoreactivity (arrowheads; high magnification in inset B-b; E-a,b); this staining pattern was also found in the AD (C). MyD88 also around the endothelial cells in blood vessels (*bv*). Tissue edema (parenchymal distension/”gaps”) is prominent (B, E). Images taken with a 40x objective. On the right panel, semiquantitative analysis of DAB images using the *ImageJ Fiji* software are shown. Data represents the mean of 8 images/fields per section ± S.E.M and are expressed as a percentage of change versus the control group. Different from control: *p < 0.05, **p < 0.01, ****p < 0.0001; different from WE: ##p < 0.01.

**FIGURE 3.**
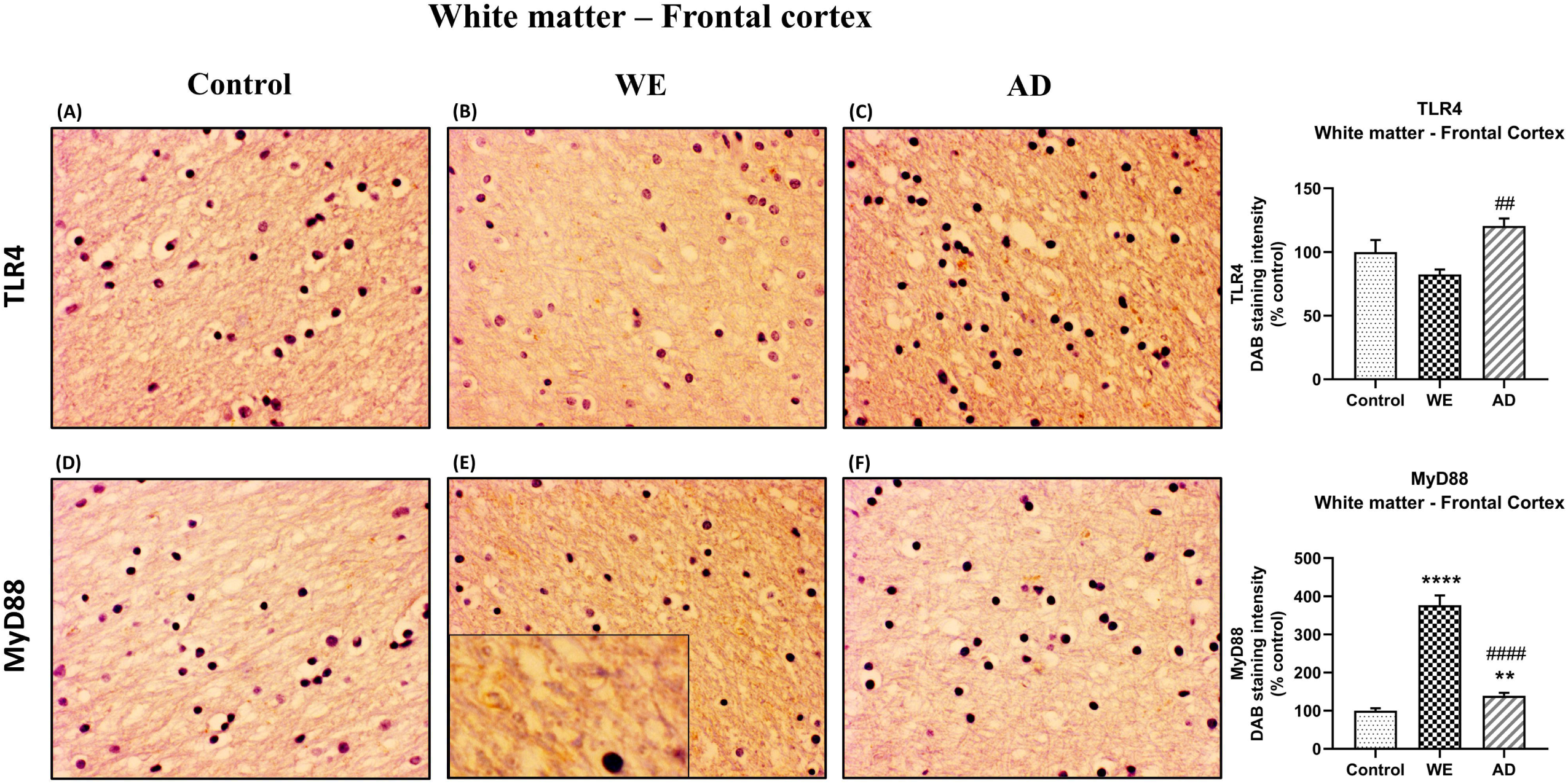
Representative images of immunohistochemical detection for TLR4 and MyD88 in the white matter of the frontal cortex of control, alcohol-related Wernicke’s encephalopathy (WE) and Alzheimer’s disease (AD) cases. WE patient showed the highest elevation in MyD88 expression (E). Images taken with a 40x objective. On the right panel, semiquantitative analysis of DAB images using the *ImageJ Fiji* software are shown. Data represents the mean of 8 images/fields per section ± S.E.M and are expressed as a percentage of change versus the control group. Different from control: **p < 0.01, ****p < 0.0001; different from WE: ##p < 0.01, ####p < 0.0001.

In the cortical gray matter of the control case, weak TLR4 immunoreactivity was occasionally detected in a few pyramidal neurons and glial cells (Figure 2A). In the same brain area of the WE patient, we observed a strong TLR4 expression in most pyramidal neurons (with cytoplasmic localization), as well as in glial cells and slightly in the neuropil. Tissue edema was also evident, with parenchymal distension (“gaps or empty spaces”) (Figure 2B). Some endothelial cells in the blood vessels also appeared to be TLR4 positive (Figure 9). Likewise, in the AD case, we also found a heavy TLR4 expression in the citoplasm of the pyramidal neurons, glial cells and in the vicinity of some blood vessels (Figures 2C and 9). Thus, we found significant differences between the cases (F _(2, 21)_ =6.758, p=0.0054), with an increased TLR4 signal in WE (p=0.0066) and AD (p=0.0358) cortical gray matter compared with the TLR4 positive staining in the control case. In addition, we found higher MyD88 expression in WE (p=0.0086) and AD (p<0.0001) compared with the MyD88 staining in the control case (Figure 2E, F; F _(2, 20)_ =24.62, p<0.0001). MyD88 immunoreactivity was observed in the cytoplasm of pyramidal neurons, in some glial cells and around the endothelial cells in blood vessels in the WE tissue (Figures 2E and 9). A similar pattern of greater intensity was observed in the AD case (Figures 2F and 9, p=0.0039).

Regarding the results found in the cortical white matter, control and WE cases showed a faint TLR4 staining (Figure 3A, B) but a higher TLR4 immunoreactivity was detected in the AD patient mostly between the fibres and in glial cells, showing the AD case a higher TLR4 expression compared to the WE case (p=0.0028) (Figure 3C; F _(2, 20)_ =7.53, p=0.0036).

Notably, we found an increase in MyD88 expression in the cortical white matter of both the WE and AD patients when compared to the control case (p <0.0001 and p=0.0018, respectively), and such an increase was particularly prominent in the WE case (p <0.0001, compared to the AD case) (Figure 3D, E, F; F _(2, 21)_ =139.4, p<0.0001). This pronounced WE positive signal appears to be detected by the surrounding fibers and glial cells (Figure 3E, magnified box).

In addition, as well as in the animals, we also checked the p-NFκB p65 and IκB-α markers. In the cortical gray matter, we noticed comparatively elevated immunoreactivity of p-NFκB p65 in the WE case versus control case (p=0.0204), finding mostly a cell nuclear localization of this mediator of inflammation. This can be observed mainly in neurons (especially pyramidal) (Figure 4B). A very similar staining pattern was also found in the positive control for neuroinflammation, the AD case, being significantly different from the control (p=0.0003) (Figure 4C) (Figure 4, p-NFκB p65 U= 16.14, p=0.0003).

Regarding to the IκB-α, in the gray matter of the frontal cortex, we found certain differences between the cases (Figure 4, F _(2, 21)_ =4.406, p=0.0253). In the WE patient, a slight staining of the cell cytoplasm was observed in some neurons, although it was not significant compared to the control (Figure 4E, p>0.05, n.s.). Similarly, the AD-positive control showed no differences in IκB-α staining versus the control (Figure 4E, p>0.05, n.s.). However, although lower total levels of immunoreactivity were detected in the AD subject compared to the WE case (p= 0.0221), it is noteworthy to highlight that a striking IκB-α labeling was observed in the astrocytes (Figure 4F-e, f), which was also found in the WE case with less intensity, with some astrocytes reacting in the same way to this marker (Figure 4E-d).

**FIGURE 4.**
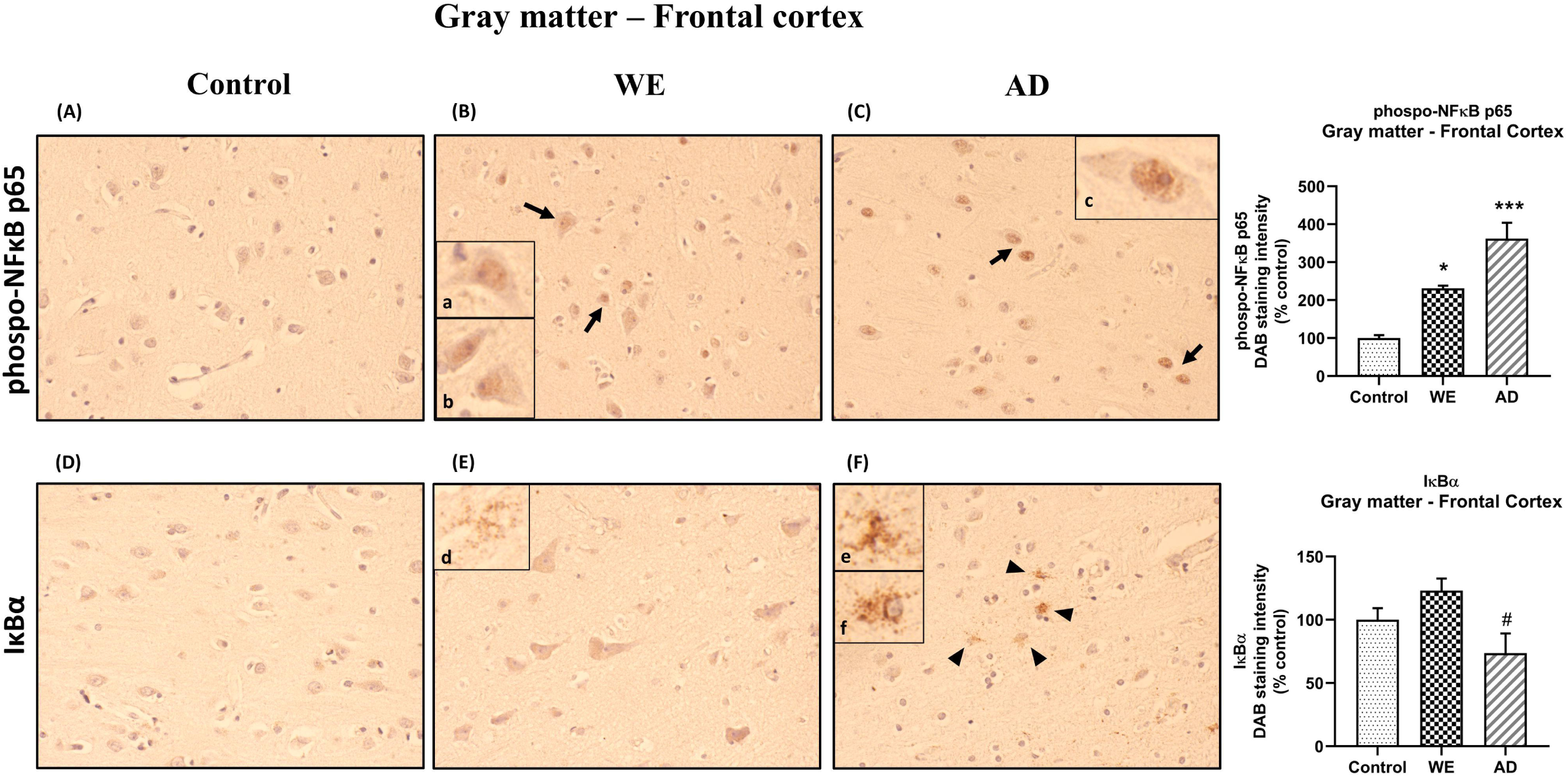
Representative images of immunohistochemical detection for p-NFκB p65 and IκB-α in the gray matter of the frontal cortex of control, alcohol-related Wernicke’s encephalopathy (WE) and Alzheimer’s disease (AD) cases. p-NF-κB exhibited nucleus localization in WE and AD cases (see especially in pyramidal neurons: arrows; high magnification in inset B-a, b; C-c). IκB-α manifested cytoplasmic localization, but highlighting a striking immunoreactivity in the astrocytes, mainly in the AD (arrowheads; high magnification in inset F-e, f), but also, although to a lesser extent, found in the WE case (high magnification in inset E-d). Images taken with a 40x objective. On the right panel, semiquantitative analysis of DAB images using the *ImageJ Fiji* software are shown. Data represent the mean of 8 images/fields per section ± S.E.M and are expressed as a percentage of change versus the control group. Different from control: *p < 0.05, ***p < 0.001; different from WE: #p < 0.05.

In the cortical white matter, we found no significant differences between cases with p-NF-κB and those with IκB-α (Figure 5, p>0.05, n.s.).

**FIGURE 5.**
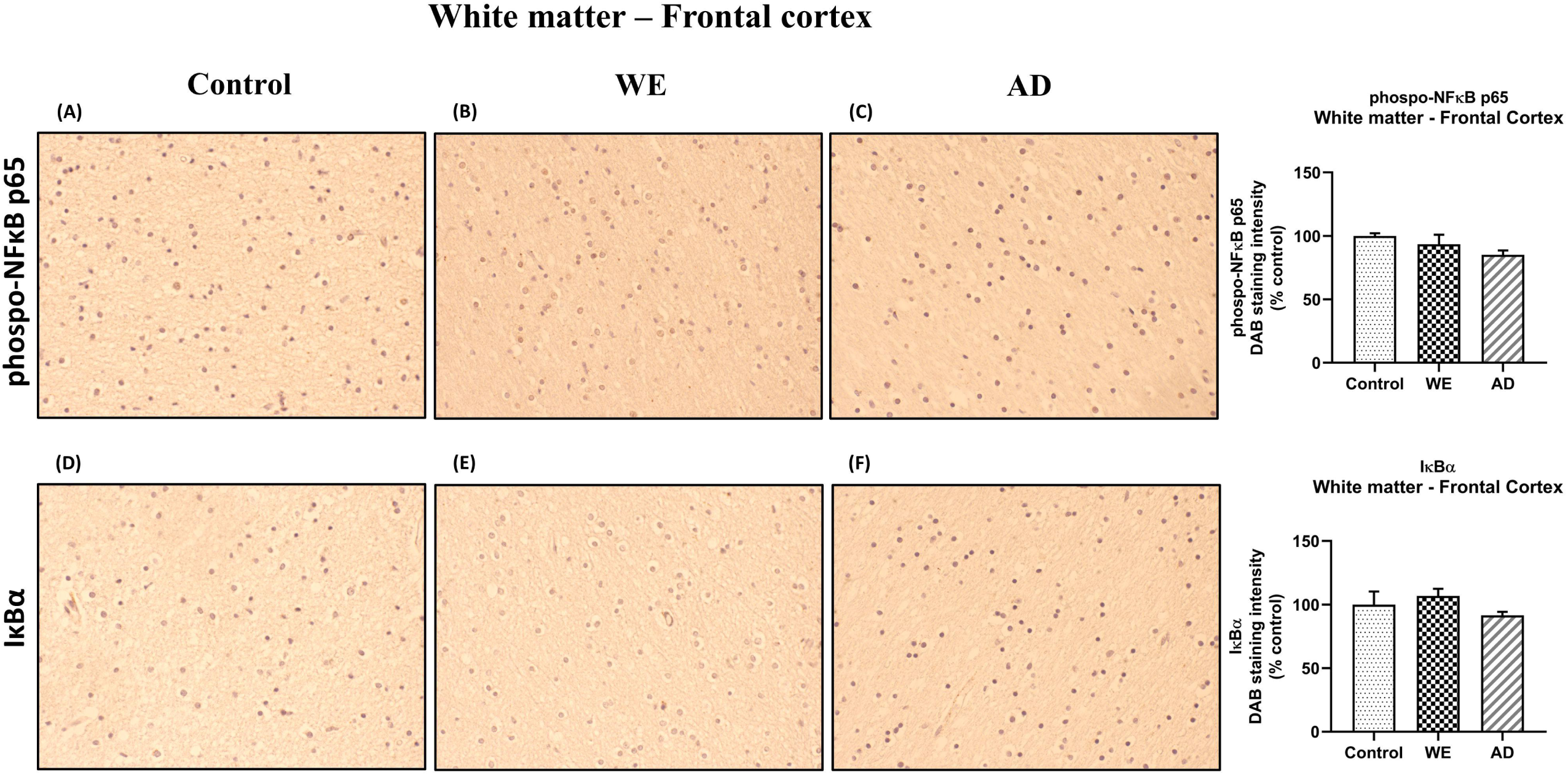
Representative images of immunohistochemical detection for p-NFκB p65 and IκB-α in the white matter of the frontal cortex of control, alcohol-related Wernicke’s encephalopathy (WE) and Alzheimer’s disease (AD) cases. Images taken with a 40x objective. On the right panel, semiquantitative analysis of DAB images using the *ImageJ Fiji* software are shown. Data represent the mean of 8 images/fields per section ± S.E.M and are expressed as a percentage of change versus the control group.

### Cerebellar Findings

#### Unaffected expression of TLR4 signaling markers in the cerebellar hemisphere of TDD, CA and CA+TDD-treated rats

None of the experimental conditions induced significant changes in the markers studied (Figure 6A, B, C, D; total NF-κB: Supplementary Data, section 2.3, Figure 5, p>0.05, n.s.). Inducible oxide nitric synthase (iNOS) enzyme and heat shock protein 70 (HSP70) were also analyzed, showing no alterations in their levels by any of the treatments (p>0.05, n.s., Supplementary Data, section 2.3, Figure 4B, C, respectively).

**FIGURE 6.**
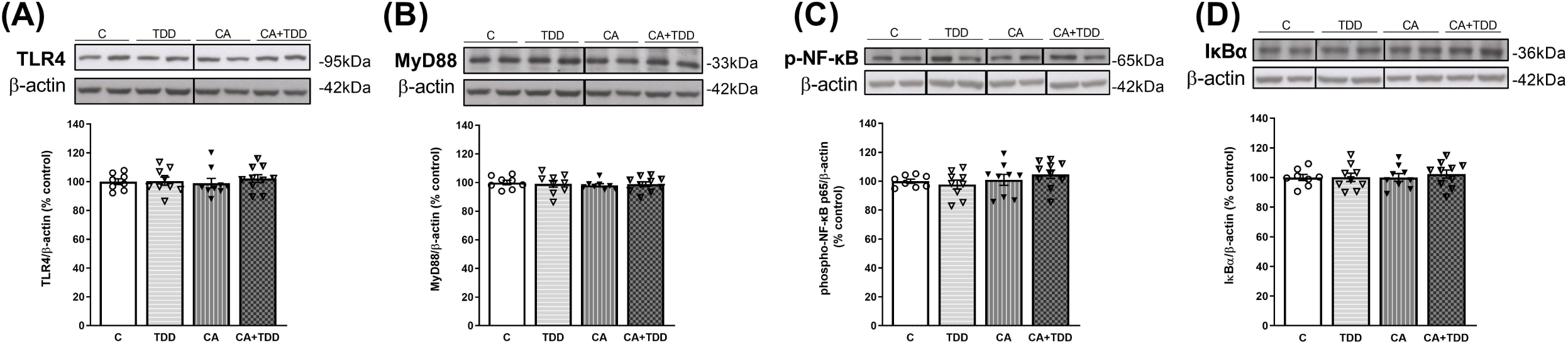
Unaffected expression of TLR4 signaling pathway in cerebellar hemisphere of TDD, CA and CA+TDD-treated rats. Graphs indicate protein levels of (A) TLR4, (B) MyD88, (C) p-NF-ĸB and (D) IĸBα markers by Western blot; data of the respective bands of interest (upper bands) were normalized by β-actin (lower band) and expressed as a percentage of change versus the control group. Some blots were cropped from the original (black lines) for improving the clarity and conciseness of the presentation. Mean ± SEM (n = 8-10). Two-way ANOVA. Since the combined CA+TDD treatment mimics better the human case of alcohol-induced WE, this group was also compared with the C group by Unpaired Student’s t-test or Mann-Whitney (CA+TDD vs C).

Likewise, no significant differences were found when comparing the CA+TDD group with the control in any of these markers (p>0.05, n.s.).

#### Postmortem human cerebellar hemisphere of alcohol-induced WE showed an increased expression of TLR4 and its co-receptor MyD88 and IκB-α immunoreactivity

In the cerebellum sections, the three cellular layers of the cerebellar cortex - the molecular layer, Purkinje cells and the granular layer – were observed and analysed together as cerebellar grey matter. The control case showed only occasional and low TLR4 immunoreactivity, mainly in some cells in the transition between the molecular and the granular layer (Figure 7A). In contrast, the WE case showed a more intense TLR4 staining, especially in the granular layer, in the cells and between the branching or neuropil; endothelial cells of blood vessels also showed TLR4 staining (Figures 7B and 9). Likewise, TLR4 in the AD patient was found mostly throughout the granular layer and in blood vessels (Figures 7C and 9). Accordingly, the semiquantitative analysis demonstrated a significant increase in TLR4 expression in the cerebellar hemisphere gray matter (F _(2, 20)_ =13.81, p=0.0002) of WE (p=0.0006) and AD (p=0.0006) patients compared to the control case.

**FIGURE 7.**
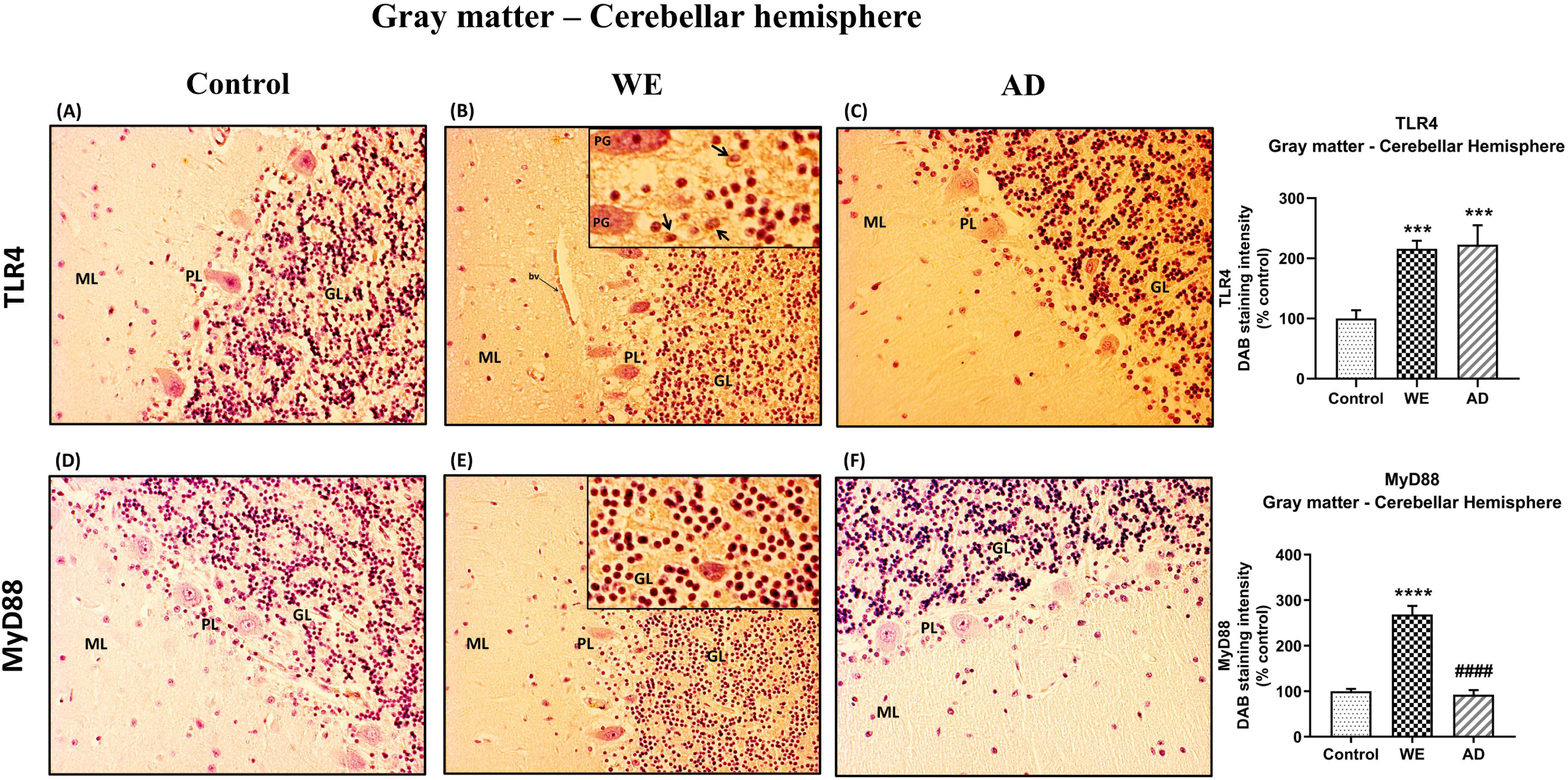
Representative images of immunohistochemical detection for TLR4 and MyD88 in the gray matter of the cerebellar hemisphere of control, alcohol-related Wernicke’s encephalopathy (WE) and Alzheimer’s disease (AD) cases. Gray matter with molecular layer (*ML*), Purkinje cell layer (*PL*) and granular layer (*GL*). In the WE (B) and AD cases (C), an increased TLR4 immunoreactivity was detected specially in the granular layer (high magnification in inset in B; Purkinje cell, PG; arrows pointing to positive cells) and blood vessels (*bv*), compared with the control (A). WE patient also showed the highest elevation in MyD88 expression, mainly by the granular layer (high magnification in inset in E), compared with AD (F) and control (D). Images taken with a 20x objective. On the right panel, semiquantitative analysis of DAB images using the *ImageJ Fiji* software are shown. Data represents the mean of 8 images/fields per section ± S.E.M and are expressed as a percentage of change versus the control group. Different from control: ***p < 0.001; ****p < 0.0001; different from WE: ####p < 0.0001.

In the cerebellar cortex, MyD88 staining was predominant in the WE patient, with a main distribution within the granular layer, as well as TLR4 (Figure 7E), whereas a weak MyD88 immunoreactivity was found in both the control case and the AD patient (Figure 7D and F, respectively). MyD88 expression intensity was significantly increased in the cerebellar hemisphere gray matter (F _(2, 21)_ =54.03, p<0.0001), showing the WE patient the highest levels compared to the AD patient (p<0.0001) and control case (p<0.0001).

In contrast, cerebellar hemisphere white matter did not show any differences in either TLR4 nor MyD88 immunostaining in the human cases analyzed (Figure 8).

**FIGURE 8.**
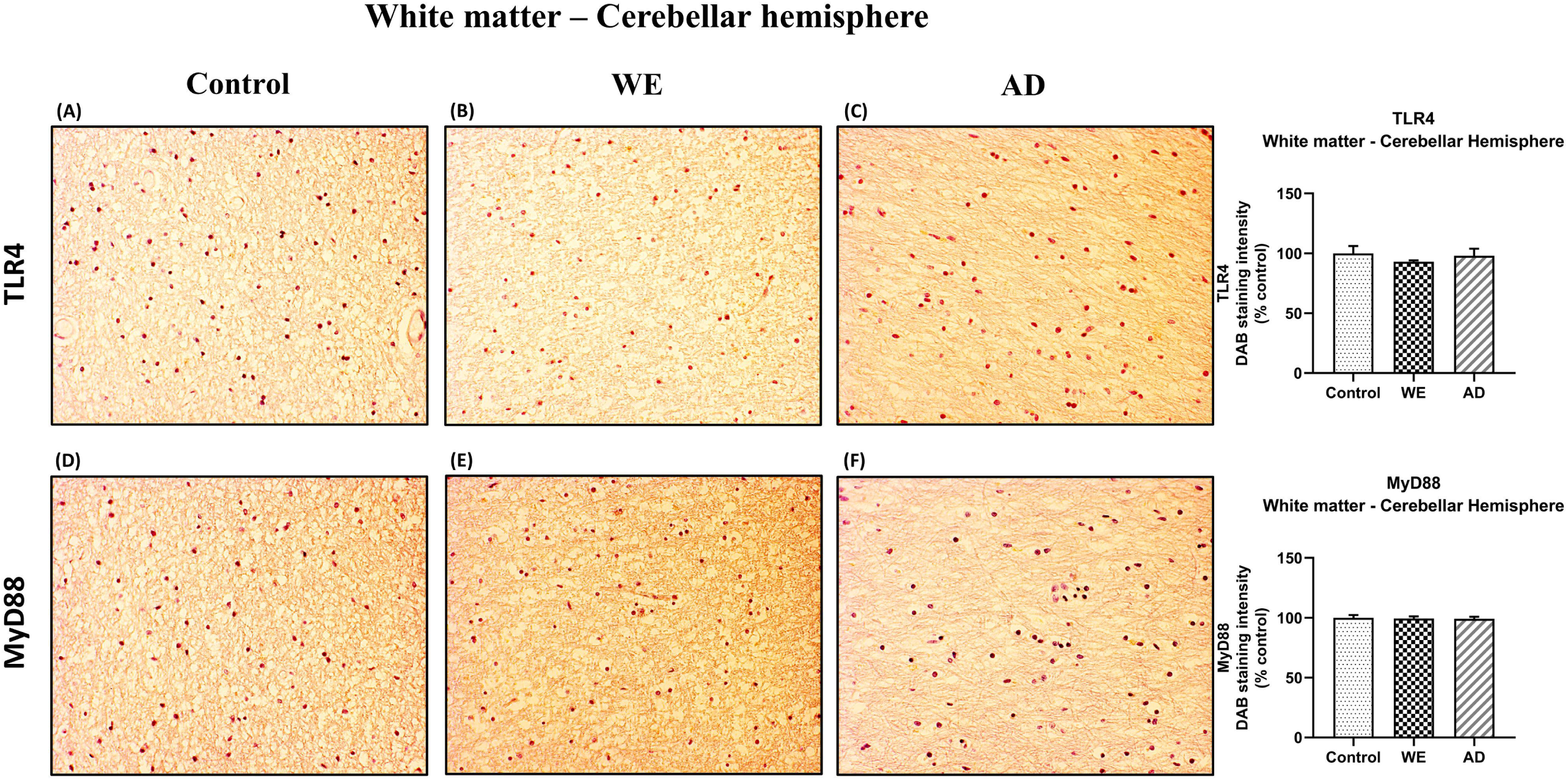
Representative images of immunohistochemical detection for TLR4 and MyD88 in the white matter of the cerebellar hemisphere of control, alcohol-related Wernicke’s encephalopathy (WE) and Alzheimer’s disease (AD) cases. Images taken with a 20x objective. On the right panel, semiquantitative analysis of DAB images using the *ImageJ Fiji* software are shown. Data represents the mean of 8 images/fields per section ± S.E.M and are expressed as a percentage of change versus the control group.

Additionally, p-NF-κB and IκB-α immunoreactivity were also analyzed in the cerebellar hemisphere, and the p-NF-κB results in the gray matter showed no differences in the WE compared to control (Figure 10, p>0,05 n.s.). There was a slight difference between the cases (Figure 10, F _(2, 20)_ =12.27, p=0.0003), with an apparent lower level of labeling in the AD case compared to the control (p=0.0002) and to the WE subject (p=0.0369).

**FIGURE 9.**
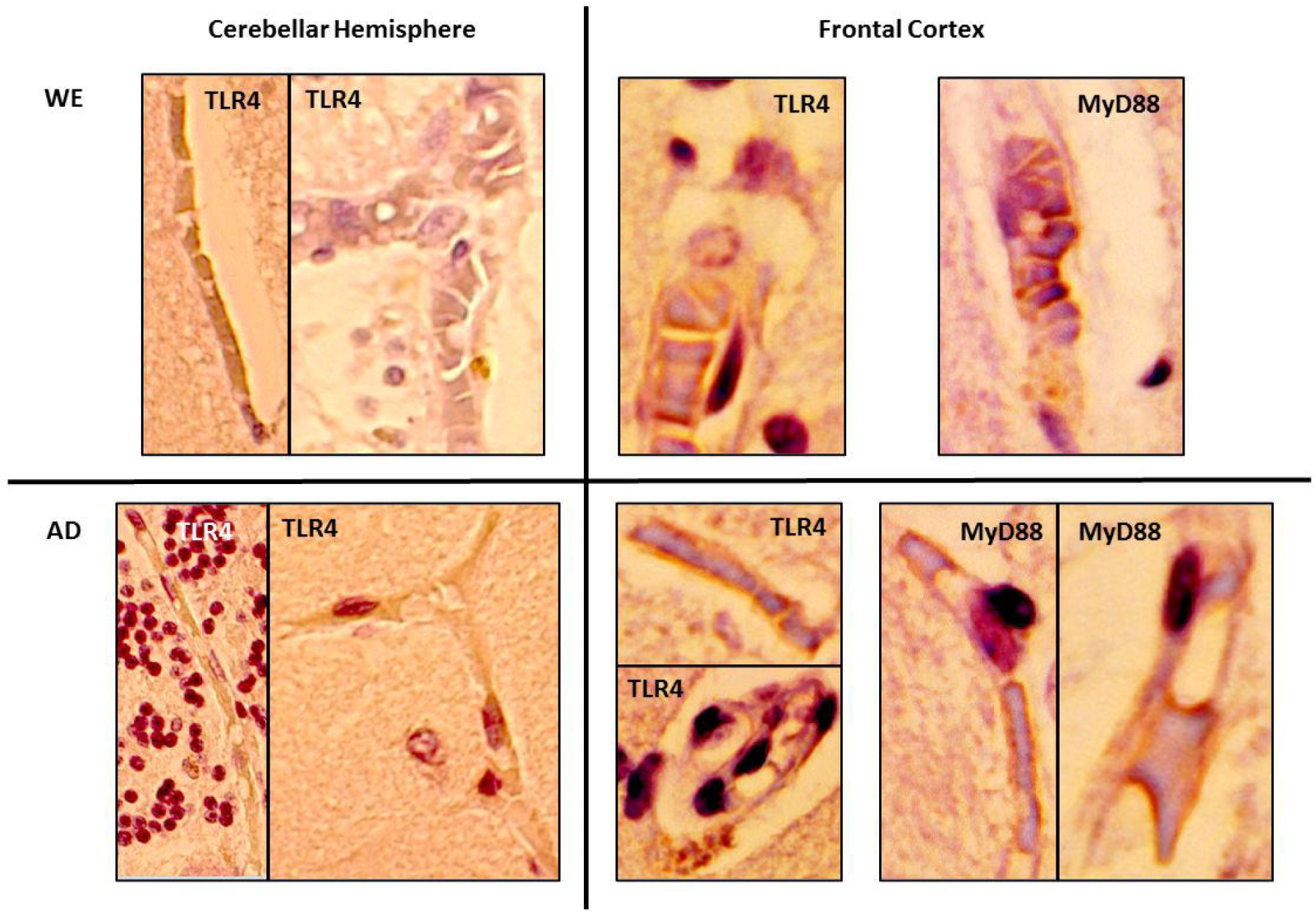
Detail showing the endothelial cells of blood vessels with immunohistochemical reactivity of TLR4 and MyD88 in the gray matter of cerebellar hemisphere and frontal cortex of alcohol-related Wernicke’s encephalopathy (WE) and Alzheimer’s disease (AD) cases. High magnification from 20x (cerebellar hemisphere) and 40x (frontal cortex) images.

**FIGURE 10.**
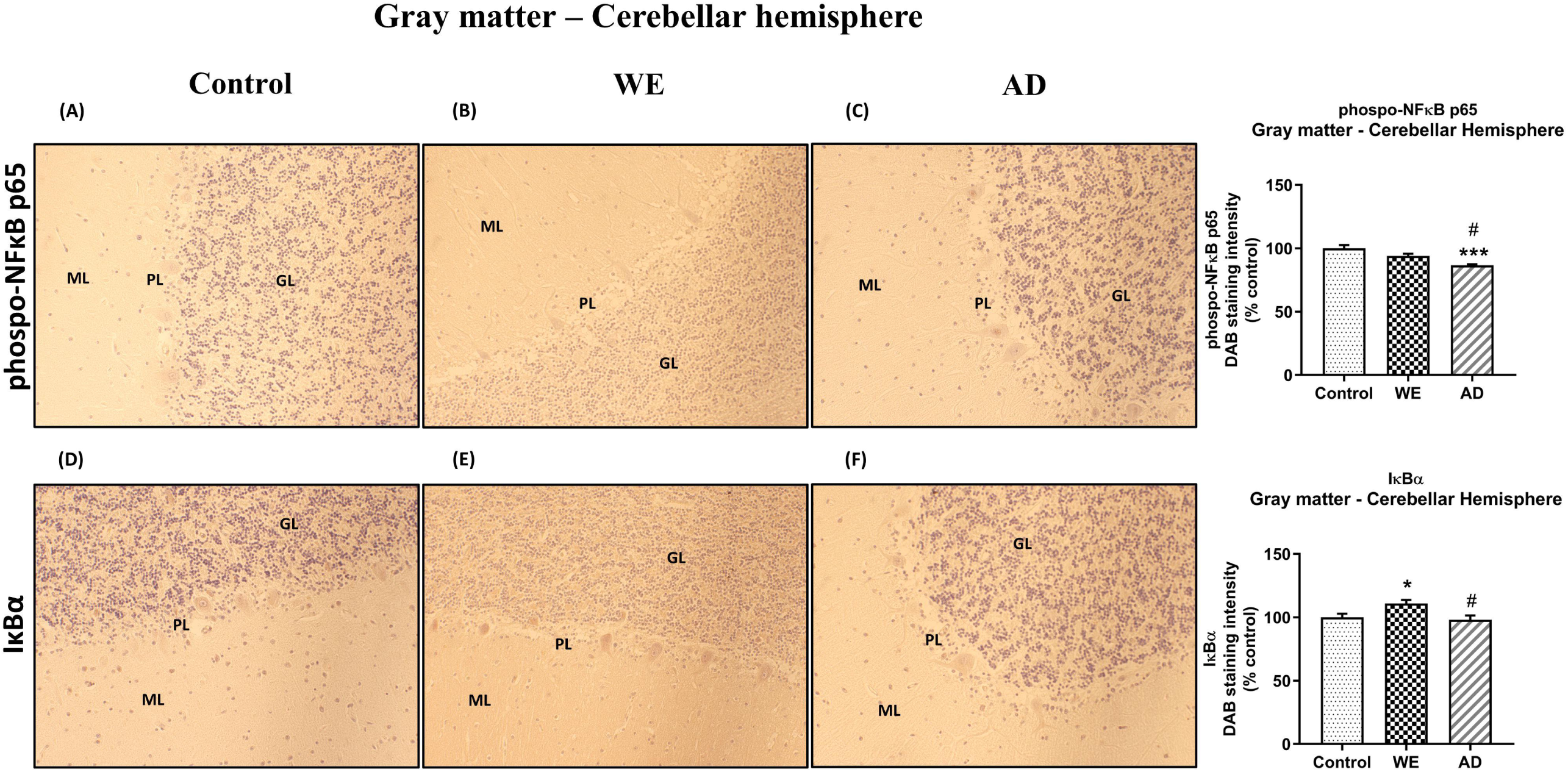
Representative images of immunohistochemical detection for p-NFκB p65 and IκB-α in the gray matter of the cerebellar hemisphere of control, alcohol-related Wernicke’s encephalopathy (WE) and Alzheimer’s disease (AD) cases. Gray matter with molecular layer (*ML*), Purkinje cell layer (*PL*) and granular layer (*GL*). In the WE (E), an increased IκB-α immunoreactivity was detected specially in the granular layer compared with the control (D). Images taken with a 20x objective. On the right panel, semiquantitative analysis of DAB images using the *ImageJ Fiji* software are shown. Data represent the mean of 8 images/fields per section ± S.E.M and are expressed as a percentage of change versus the control group. Different from control: *p < 0.05; ***p < 0.001; different from WE: #p < 0.05.

With regard to IκB-α, we observed significant differences between the patients (Figure 10, F _(2, 20)_ =5.466, p=0.0128), finding an increased IκB-α immunoreactivity in the WE case compared to control (p=0.0462) and to the AD subject (p=0.0207). This IκB-α-labeling appears to be observed mainly through the granular layer, as occurred with TLR4 and MyD88.

In agreement with the results of TLR4 and MyD88 found in the white matter of the cerebellar hemisphere, we observed no significant changes between cases in this area in p-NF-κB and IκB-α markers (Figure 11, p>0.05, n.s.).

**FIGURE 11.**
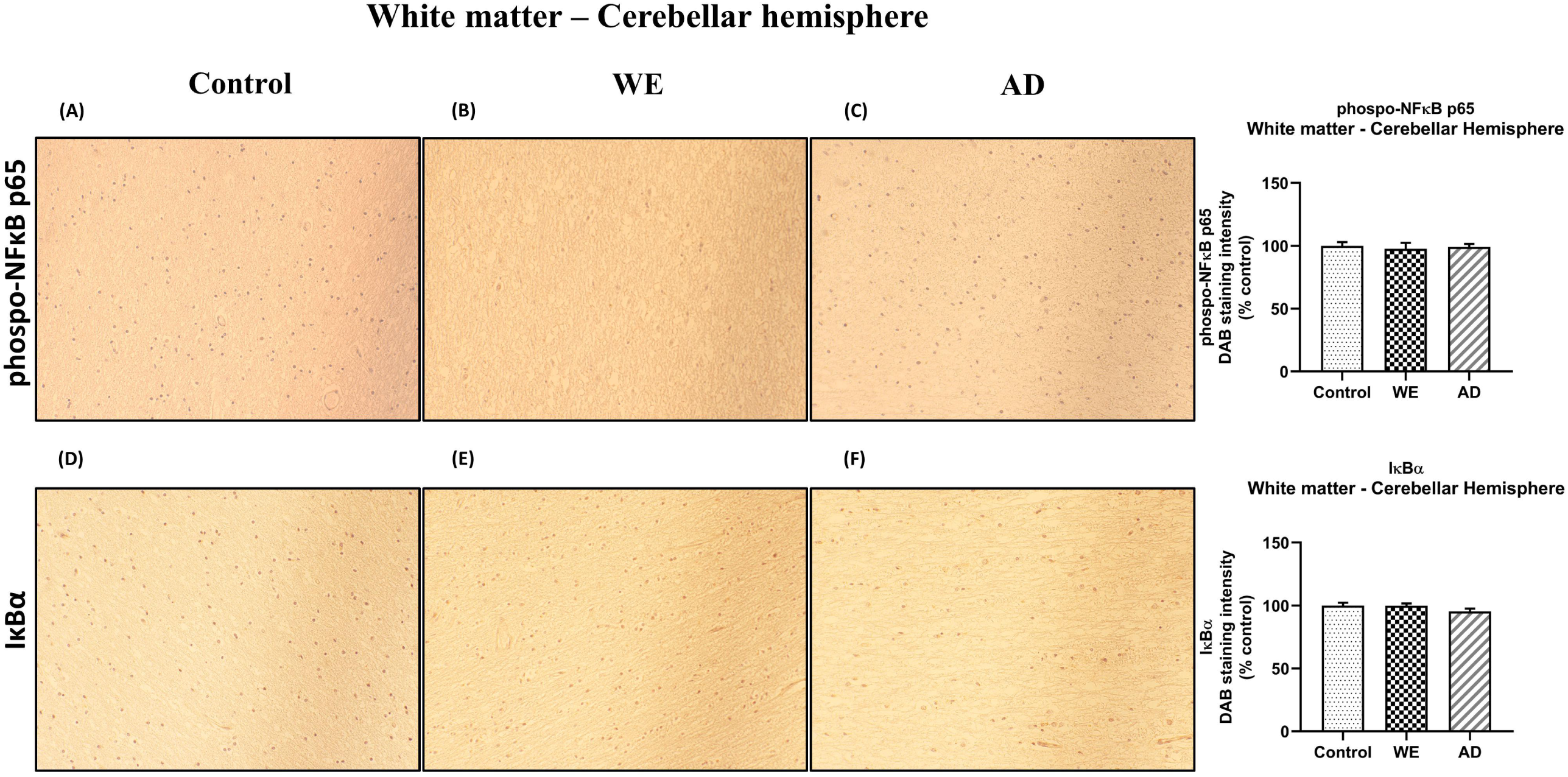
Representative images of immunohistochemical detection for p-NFκB p65 and IκB-α in the white matter of the cerebellar hemisphere of control, alcohol-related Wernicke’s encephalopathy (WE) and Alzheimer’s disease (AD) cases. Images taken with a 20x objective. On the right panel, semiquantitative analysis of DAB images using the *ImageJ Fiji* software are shown. Data represent the mean of 8 images/fields per section ± S.E.M and are expressed as a percentage of change versus the control group.

#### Thiamine levels

Plasma thiamine levels were measured in all animals of our study. Briefly, after 9 months of exposure to alcohol and after TD diet treatment, we found a trend towards a decrease in total thiamine levels due to an alcohol effect (for detailed results see (Moya et al., 2022).

In the case of the WE patient studied here, it was not possible to perform thiamine determinations, since she died very quickly. Nevertheless, neuropathological analyses confirmed the diagnosis of WE without comorbidity with other pathologies such as HE, as we have already explained.

#### Liver status

We checked the status of the liver in the animals by measuring the hepatic nitrites MDA levels, due to the major role of these processes in the pathogenesis of ALD (McKim et al., 2003; Galicia-Moreno and Gutiérrez-Reyes, 2014; Pérez-Hernández et al., 2017; Tan et al., 2020; Yang et al., 2022). The levels of hepatic MDA and NO_2-_ showed no significant changes by any of the treatments (Supplementary Data, section 2.4, Figure 6).

Regarding the human case, the liver enzymes’ (transaminases) values were in the upper limit, but without exceeding reference levels (See Supplemental Data, section 1.1). The patient exhibited no other symptoms or clinical signs of liver disease, suggesting that she was an alcohol-induced WE patient without alcoholic liver disease (ALD).

## DISCUSSION

Little is known about a possible role of TLR4 in the WE, since there are no studies in humans and, to our knowledge, only our previous work with TD-animal models provides evidence about the contribution of this receptor to this pathology (Moya et al., 2021). In the present study, we further characterized the role of this receptor in WE by studying animal models with combined TD and chronic alcohol use. Thus, here we report the importance of the double hit (TD and chronic alcohol use) in the magnitude of the expression of the proinflammatory TLR4 signalling cascade in the frontal cortex but not in the cerebellum. We also described the presence of an upregulated cortical and cerebellar TLR4 and its adaptor molecule MyD88, along with specific changes in the signaling molecules phospo-NFκB p65 and IκBα, in a single case of alcohol-induced WE, by using postmortem brain tissue.

WE patients show neuropsychological symptoms such as memory alterations, apathy, executive deficit and disinhibition, which suggest a dysfunction of frontal structures. The vulnerability of the frontal lobe to chronic alcohol consumption with or without TD is widely accepted based on neuropathological and neuroimaging studies (reviewed in (Jung et al., 2012). Inflammation, among other processes, may contribute to the WE symptomatology, as it increases cell damage and causes neuronal death. The innate immune receptor TLR4 play a critical role in determining the pathological outcomes in several neurological and neuropsychiatric disorders, including AUD, AD, depression, schizophrenia or trauma (Crews et al., 2013; García Bueno et al., 2016). By using WE animal models resulting from TD exposure, we were able to identify TLR4 as a key molecule in the emotional, cognitive and motor disturbances associated to these models in a previous study (Moya et al., 2021). To our knowledge, there are no previous works that examine TLR4 signalling in the brain in the context of WE, either in animals or in human subjects.

However, since the main documented cause of WE is alcohol consumption, it is needed to explore more complex animal models in which we can combine TD and chronic alcohol consumption and explore the specific contribution of each factor to the pathophysiology of the disease. Indeed, in a very recent publication of our group we described the contribution of both factors (TD and alcohol abuse) to the induction of neuronal damage in the frontal cortex and how the combination of both (CA+TDD) correlates with disinhibition-like behavior in animals (Moya et al., 2022), which is a core symptom of the pathology. Similarly, in the present study, we used the same combined animal models to explore the specific role of TLR4 in the induction of a neuroinflammatory cascade in the frontal cortex and cerebellum, and added the description of the TLR4/MyD88 upregulation in postmortem brain of an alcohol-induced WE case.

Among the animal models investigated here, the CA+TDD model is the one that better represents the pathology, as expected (Moya et al., 2022) and it is also the best approach to be compared with the alcohol-related WE case (postmortem human brain) studied here. In the animal model, we observed a significant up-regulation in the protein expression of TLR4, MyD88, p-NF-κB and IĸBα in the frontal cortex (CA+TDD versus control group). It is to note that the neuroinflammation is a very complex response which involves the activation of several factors. The upregulation of phosphorylated NF-κB p65 is indicative of activation of this nuclear factor, since phosphorylation of Ser536 in the cytosolic p65 promotes the nuclear translocation and facilitates p65 binding to specific promoter sequences, activating the inflammatory gene expression (Giridharan and Srinivasan, 2018). Additionally, its inhibitor gene IκBα contains NF-κB binding sites in its promoter, so the NF-κB is able to autoregulate the transcription of this own inhibitor, meaning that the “NF-κB-IκBα autoregulatory feedback loop” would be trying to suppress a prolonged activation of NF-ĸB to limit the inflammatory response (Doremus-Fitzwater et al., 2015; Gano et al., 2016; Toledo Nunes et al., 2019; Moya et al., 2021). Indeed, this autoregulatory mechanism switched on by NF-κB to block its stimulation is widely known in neuroinflammatory studies induced by LPS (Sayd et al., 2015), induced by alcohol (Doremus-Fitzwater et al., 2015; Gano et al., 2016), by TD (Moya et al., 2021) or combined TD and alcohol (Toledo Nunes et al., 2019). So, the IĸBα protein levels are useful reporters of NF-ĸB activity and increase neuroinflammatory status.

It could be surprising that the TLR4 was not upregulated in the TD model, as opposed to our previous studies (Moya et al., 2021). It is known that neuroinflammation is a complex response where markers peak at different time-points, so the lack of significant effect in TLR4 in the TD animals of this study could be indicative that we did not catch the TLR4 peak at the precise moment of the samples collection. This is probably related with the age difference between the animals in both studies. Whereas in the previous study the animals were 8-9 weeks old, in the current study they were approximately 10 months old. It is very possible that the younger animals react differently (they show a particular timing of TLR4 up-regulation profile) than the older animals. Nevertheless, there is certain evidence of a TLR4 signaling pathway overactivation in these animals, as some other inflammatory signals were observed (MyD88, IκBα) in the frontal cortex. Indeed, the upregulation of the TLR4 co-receptor MyD88 could be interpreted as a sign of TLR4 signaling overresponse, in absence of significant receptor overexpression. However, TLR4 is upregulated in the combined animal model, CA + TDD, although the increase is moderate compared to controls. In this regard, it is to note that the animals were exposed to a chronic treatment of moderate alcohol intake, so we are facing a process of chronic neuroinflammation, where we cannot expect such pronounced elevations as in a *binge drinking* model for example, where there is a peak of acute neuroinflammation with a prominent increase in cortical levels of the TLR4-pathway (Antón et al., 2017). Additionally, there is an absence of synergic effect by the combination of CA and TDD, since the elevations in the TLR4 pathway proteins were not higher than in the CA and/or TD exposure alone, which is maybe a consequence of the long alcohol exposure (9 months), rendering cells less sensitive to the TDD response.

It is to note that, together with the upregulation of TLR4 neuroinflammatory pathway found in CA+TDD animals this study, we have already described, to complement the mechanisms, that other processes associated to TLR4 activation such as oxidative and nitrosative stress, lipid peroxidation, apoptosis death and cell damage are upregulated in the frontal cortex of the same animals and correlate with disinhibition-like behavior (Moya et al., 2022). All these markers are more representative of the latest stages of a neuroinflammatory response, traditionally linked to neurotoxicity (Moya et al., 2022). So, altogether, these studies shed light on the relative contributions of each factor (alcohol and TD, either isolated or in interaction) to the potential disease-specific mechanisms involved in the WKS pathophysiology, resulting in brain damage and behavioral problems.

In a complementary way, here we show, for the first time, a case of WE associated to chronic alcohol consumption in which there is an upregulation of TLR4 and MyD88 protein expression in the postmortem frontal cortex and cerebellum. Neuroinflammation involves all the cell types present within the central nervous system (Shabab et al., 2017). In this way, microglia and astrocytes, as well as neurons and oligodendrocytes all contribute to innate immune responses in the CNS through the expression of TLR4, among other TLRs. Thus, TLR4 is expressed in human brain cells, including neurons, microglia, astrocytes and oligodendrocytes (Vaure and Liu, 2014; Stephenson et al., 2018; Frederiksen et al., 2019; Kumar, 2019; Leitner et al., 2019). In the WE cortical gray matter, immunohistochemical analysis showed an increased TLR4 staining in glial cells and pyramidal neurons, mainly in the cytoplasm, since TLR4 can signal both at the plasma membrane and intracellularly (Gangloff, 2012). TLR4 immunoreactivity was also observed in endothelial cells of the blood vessels, in agreement with Nagyoszi and colleagues, who demonstrated the expression of TLR4 on rat and human cerebral endothelial cells induced by inflammatory stimuli or oxidative stress (Nagyoszi et al., 2010). MyD88 expression in the WE patient showed a staining pattern similar to that observed for TLR4: it was mainly detected in pyramidal neurons and glial cells. Such a result may suggest that the upregulation of TLR4 has functional consequences in the associated signaling pathway. Likewise, endothelial cells of blood vessels showed immunoreactivity, which could fit with a study reporting that activation of MyD88 pathway in endothelial cells of the cerebral microvasculature is involved in the regulation of the inflammatory events (Gosselin and Rivest, 2008).

Similar findings have been previously reported in the AD brain, considered as a positive control, showing an activation of TLR4 in both human AD diagnosed patients and AD animal models (Fiebich et al., 2018; Calvo-Rodriguez et al., 2020; Zhou et al., 2020). The increase in TLR4 expression was particularly observed in the frontal cortex of AD subjects when compared to age-matched controls (Miron et al., 2018). Accordingly, MyD88 levels were also reported to be elevated in the cortex of patients with AD and in a mouse model of AD (Rangasamy et al., 2018).

Preclinical and human studies have demonstrated that exposure to severe alcohol alone or combined with TD leads to white matter damage in the cortex (Kril et al., 1997; Harper, 2009; de la Monte and Kril, 2014; Chatterton et al., 2020) suggesting that neuroinflammation participates in the myelin and white matter disruptions (Alfonso-Loeches et al., 2012; Toledo Nunes et al., 2019). Here, we found a prominent increase in MyD88 immunoreactivity in the WE cortical white matter, and this excessive signalling could be leading to lower TLR4 levels in this cortical area by a compensatory downregulation mechanism. Indeed, depending on the temporal status in which these parameters where measured, the balance between TLR4 and MyD88 upregulation can be differentially affected.

Thereby, the concurrence of TLR4 and MyD88 immunoreactivity suggests an activation of the TLR4-MyD88 signaling pathway, although we cannot exclude other alternative pathways. It is possible that other TLR4 MyD88-independent signal transduction pathways such as the TRIF-dependent pathway could also be activated. So, the TLR4 immunoreactivity detected in our study could be indicative of signaling from both the membrane through Myd88 and internally from TRIF, as well as the MyD88 immunoreactivity could be somehow non-specific for TLR4 and may include other TLRs (Biswas, 2018). Nevertheless, even if different pathways are activated, all of them converge and activate the NF-κB factor, in which we are particularly interested because it is the foremost important transcriptional manager of inflammation-associated genes (Marongiu et al., 2019; Ciesielska et al., 2021; Lin et al., 2021; Duan et al., 2022).

Indeed, we found an increase of the proinflammatory mediator p-NF-κB in the cortical gray matter of the WE case compared to control, as well as in the positive-AD-control, with a predominant expression or nuclear localization, indicating that this proinflammatory factor is active, which is, presumably, a direct consequence of the activation of the TLR4 signaling in the frontal cortex. Results regarding IκB-α are sometimes difficult to explain as both factors regulate their levels through compensatory mechanisms, as explained above. In this study we found an interesting striking pattern of IκB-α labeling in glial cells such as astrocytes in the AD case, which was reproduced, to a lesser degree, in the WE patient.

Regarding the cerebellum, a damage induced by alcohol and TD has been previously reported (Mulholland, 2006; Manzo et al., 2014). Moderate shrinkage of the vermis and cerebellar hemispheres was observed in post-mortem examination in patients diagnosed with alcohol abuse and with WE (Harper, 1979). Interestingly, our analyses in the postmortem cerebellar hemisphere of the WE patient showed an increase in TLR4 expression compared to the control brain, mainly detected in the granular layer and in endothelial cells of blood vessels. In addition, MyD88 and IκB-α were also upregulated and observed mostly in the granular layer of WE subject. Accordingly, the cerebellum is also considered as a vulnerable region for AD pathology (Hoxha et al., 2018). AD patients showed a severe astrocytosis in the cerebellar granular layer (Fukutani et al., 1996), and studies with cerebellar granule cells reported increased secretion of ß-amyloid related with the neurodegeneration of nearby cells (Galli et al., 1998) (reviewed in (Hoxha et al., 2018)). Here, we also found elevated TLR4 signal in the cerebellar hemisphere of the AD patient compared with the control case, although this increase was not observed for MyD88. As explained above, we cannot exclude the implication of other independent pathways to MyD88, so TLR4 could mediate their effects through different inflammatory mediators in this brain area in the AD.

In contrast to the data found in the WE patient, none of the treatments appeared to affect the TLR4 signalling in the cerebellar hemisphere of the rats compared with controls. In our previous study (Moya et al., 2021), the 12 days TD-induced model did not show neuroinflammation signature in the cerebellum, coincident with an absence of motor impairment. However, we found the neuroinflammatory markers increased in the cerebellum in another model with a deeper degree of TD due to severe TDD treatment of 16 days, where the decline in animals’ motor performance positively correlated with an upregulation of p65 NF-κB in this brain region (Moya et al., 2021). In the present study we observe no evidence of motor dysfunction, such as ataxia, in any of the animal models tested, thus providing an explanation for the lack of changes observed in the cerebellum. However, it is to note that the clinic history of the WE patient showed hypotonia, hyporeflexia, oculomotor deficits as nystagmus and saccadic intrusions, as well as an altered speech, which are presumably signs of cerebellar dysfunction (Bodranghien et al., 2016; Jafar-Nejad et al., 2017) and may explain the changes observed in the cerebellum of the post-mortem WE brain. Thus, the TLR4 signature in the cerebellum appears to precisely coincide with the manifestation of cerebellar symptoms, as reported by us previously (Moya et al., 2021).

Finally, it is noteworthy to take into account possible hepatic alterations when this pathology is induced by alcohol consumption. It is known that the hepatotoxic properties of alcohol abuse may lead to ALD. However, in spite of alcohol consumption is the main cause of WE, the prevalence and characteristics of the relationship between ALD and WE remain unclear due to the lack of available data (Chamorro Fernández et al., 2011). ALD is a possible comorbidity in WE patients, which present specific clinical, analytical and radiological characteristics and a poorer prognosis compared to alcoholic WE patients without ALD (Novo-Veleiro et al., 2022). Hepatic encephalopathy (HE) induced by chronic alcohol consumption is the extreme example of brain and liver interaction. Although both HE and WE occurs frequently in the setting of alcoholism, HE is due to liver disease and/or shunting of portal blood around the liver resulting in altered metabolism of nitrogenous substances, whereas WE is due to a deficiency of thiamin (Schenker et al., 1980). Three types of HE are traditionally differentiated (A, B and C) (Weissenborn, 2019), but, in term, HE is caused when toxins that are normally cleared from the body by the liver accumulate in the blood, eventually traveling to the brain. Elevated levels of ammonia appear to play a central role in this disorder, primarily by acting as a neurotoxin that generates astrocyte swelling, resulting in cerebral edema and intracranial hypertension. Other factors, such as oxidative stress, neurosteroids, systemic inflammation, increased bile acids, impaired lactate metabolism, and altered blood-brain barrier permeability likely contribute to the process of HE (Liere et al., 2017; Hadjihambi et al., 2018). In patients with underlying liver cirrhosis, distinguishing between HE and WE sometimes becomes a tough problem (Novo-Veleiro et al., 2022). HE is characterized by a wide spectrum of nonspecific neurological, psychiatric and motor disturbances, so most of them may coincide with those in WE, since no mental alteration is unique for both disorders. Notwithstanding, mental alteration is usually a most noticeable symptom of WE (Zhao et al., 2016). The recent ISHEN (International Society for Hepatic Encephalopathy and Nitrogen Metabolism) consensus uses the onset of disorientation or asterixis as the initial sign of overt HE (Ferenci, 2017). Ultimately the diagnosis of HE is based on history and physical examination, exclusion of other causes of altered mental status, and by the laboratory clinical findings, and sometimes confirmed by a trial of therapy for this disorder. Therefore, when difficulties exist in distinguishing between HE and WE, intravenous vitamin B1 can be considered as a discriminative method or a preemptive treatment (Zhao et al., 2016).

Nevertheless, although most heavy drinkers develop fatty liver, only a 20–40% subset of patients progresses to alcoholic hepatitis, and about 10–15% develop frank cirrhosis (Ghosh Dastidar et al., 2018). The WE case studied here is an alcoholic patient with WE without ALD, since she did not develop severe liver injury, thus far from being comorbid with HE.

Likewise, regarding to our animal model of chronic alcohol consumption, the existing literature indicates that rodent models exposed to a chronic alcohol administration equivalent to the one performed in this study developed mild or moderate steatosis, but no inflammation, no fibrosis and no portal hypertension (Nevzorova et al., 2020). The steatosis or fatty liver is relatively benign and represents the initial stage in the ALD spectrum. To achieve greater damage in the liver, the alcohol drinking model is combined with other stressors to stimulate inflammation, fibrosis or hepatocellular carcinoma. These second-hit models include additional factor(s) as dietary, chemical, genetic manipulations or single or multiple alcohol binges to facilitate progression to advanced ALD (Ghosh Dastidar et al., 2018; Lamas-Paz et al., 2018; DeMorrow et al., 2021).

Notwithstanding, we checked the status of the liver in the animals by measuring the hepatic nitrites and MDA levels, due to the major role of these processes in the pathogenesis of ALD (McKim et al., 2003; Galicia-Moreno and Gutiérrez-Reyes, 2014; Pérez-Hernández et al., 2017; Tan et al., 2020; Yang et al., 2022). The results suggest that the protocol of chronic alcohol consumption used in this study did not produce oxidative damage in the liver in the long-term, since both the nitrite and the MDA levels, indicative of nitrosative stress and lipid peroxidation, respectively, showed no significant changes in the CA and CA+TDD animals versus controls. We cannot discard that the chronic alcohol consumption has produced some mild to moderate alterations in the liver of our animals, but in that case it has not apparently progressed to a state of inflammatory/oxidative injury. Thus, these results suggest that the brain inflammatory response found in this study was achieved in absence of deep liver alterations.

### Limitations, strengths, and future perspectives

Several limitations of our study should be acknowledged. On one hand, our three human cases consisted of women and only male rats were used, which does not represent the real population of the disease. Further studies are needed to investigate the potential sex differences on this TLR4 pathway. On the other hand, in the control human case, ischemic anoxia was detected in the postmortem neuropathological diagnosis. However, signs of ischemia or hypoxia were not observed in the samples employed here. Moreover, since there is evidence for an involvement of inflammatory pathways, including TLR4 upregulation, after ischemia or hypoxia (Paschon et al., 2016; Mohsin Alvi et al., 2020), the present results may suggest an even higher increase in the TLR4 inflammation pathway if the data had been compared to another healthy control. Also, there is a difference of 9 years of age between the WE patient and the control. Nevertheless, we consider it to be within a comparable valid range, as observed in other studies where the difference between controls and patients also ranges between 8-10 years (Dabos et al., 2015; Ishiki et al., 2015; Ivanski et al., 2018). In addition, we are aware that the sample size in the postmortem brain study is very limited, but due to the poor records of WE cases, the access to postmortem tissue is very complicated, so the obtained results in this study are even more noteworthy. The postmortem study is descriptive and results should be considered as a pilot study and non-firmly-conclusive. Also, we are aware that the results in humans and animals have been obtained by different methodological techniques, so in future studies we will verify these findings both by these and other methods. Nevertheless, to obtain results pointing in the same direction coming from two different techniques is also interesting and noteworthy. This indicates us that both methodologies complement each other supporting common conclusions.

Despite these limitations, this is the first study to characterize TLR4 in the frontal cortex and cerebellum of a human subject with WE. Moreover, we demonstrated the utility of the automatic *Fiji* and the visual IRS methods to analyze particularly DAB-based IHC images, since a strong correlation between both results was observed. Notwithstanding, automated analysis was chosen for the results report by reducing subjective bias and allowing to detect signal not so easily identifiable to the naked eye by the observer.

Future research is required to analyse more markers of this TLR4 signalling pathway in post-mortem cerebral tissue from WE patients, including the exploration of other vulnerable brain regions. Moreover, the study of WE cases with other aetiology, as non-alcoholic patients, is also needed.

## CONCLUSION

Taken together, our study shed light on the relative contributions of alcohol consumption and TD, either isolated or in interaction, to the activation of the TLR4/MyD88 signaling, which may act as an underlying mechanism to the pathogenesis of WE. The findings provided here using animal models, along with complementary results (Moya et al., 2022), and our previous work (Moya et al., 2021) suggest that the TLR4/MyD88 signaling may be a potential disease-specific mechanisms involved in the WE pathophysiology, resulting in brain damage and behavioral problems. We provide also first preliminary evidence of the TLR4/MyD88 upregulation in the postmortem brain tissue of a human case of WE.

Our results offer valuable information to guide future studies to further investigate these specific inflammatory mechanisms in the context of WE. The knowledge about how the inflammatory response is triggered in the WE brain and its relationship with the course of the disease is critical to understand this disabling disorder and developing new therapeutic strategies.

## DATA AVAILABILITY STATEMENT

The datasets presented in this study can be obtained by reasonable request to the authors.

## ETHICS STATEMENT

The studies involving human participants were reviewed and approved by Drug Research Ethics Committee (Comité Ético de Investigación con Medicamentos/Investigación Clínica, CEIm del HUFA, ref. 62-2018). Animal studies were approved by the Animal Welfare Committee of the Complutense University of Madrid (reference: PROEX 312-19) and conducted in compliance with the Spanish Royal Decree 118/2021 and following the European Directive 2010/63/EU on the protection of animals used for research and other scientific purposes.

## AUTHOR CONTRIBUTIONS

Study design: MM & LO; Experiments: MM, BE; Imaging: MM, EMM, MLG, EGB, CG; Data analysis: MM, EMM, MLG, LO; Original draft: MM; Revision & funding: LO. Final version revision: all authors.

