## Supplemental Information for "“Upregulation of TLR4/MyD88 pathway in alcohol-induced Wernicke’s encephalopathy: findings in preclinical models and in a postmortem human case”"

### *Supplementary Material*

#### 1 Supplementary Materials and Methods

##### 1.1. Detailed patients' medical history

**Wernicke's encephalopathy (WE) case:** A 62-year-old woman with no medical history of interest was admitted to hospital for confusional syndrome. After the studies performed the main diagnosis was WE, along with macrocytic anemia, folic acid deficiency and severe enolism. Personal and family history: she had a history of chronic alcohol consumption of at least 1 bottle of wine per day for 10-15 years, although she denied the use of toxic substances. She had no arterial hypertension, no dyslipidemia (DLP) and no diabetes (DM). It was not possible to perform thiamine determinations, since she died very quickly. The patient did not present clinically demonstrated liver pathology since there was not liver biopsy. The liver enzyme (transaminases) values are in the upper limit, but without exceeding reference levels.

| Transaminases | **WE PATIENT VALUES** | NORMAL RANGE |
|---|---|---|
| *glutamic oxaloacetic transaminase* (**GOT**) | **40 U/L** | 10-40 U/L |
| *glutamic pyruvic transaminase* (**GPT**) | **36 U/L** | 7-40 U/L |
| *gamma-glutamyl transferase* (**GGT**) | **43 U/L** | 6-50 U/L |

The patient exhibited no other symptoms or clinical signs of liver disease, so it appears that this WE case is an alcoholic patient with WE without alcoholic liver disease (ALD). Family history of dementia diagnosed as AD on maternal side. Drugs: she had Prozac treatment. No adverse drug reactions.

Neurology: spontaneous language very scarce and incoherent. She obeys simple orders. Oriented in person but disoriented in space and time. Limitation of ocular supranuclear profile supreaversion (oculocephalic reflex in the vertical plane preserved). Slowing of spontaneous eye saccades in the horizontal plane (nystagmus). Alertness, although with some tendency to drowsiness. Bradypsychia. Weak sucking reflex. Palmomenten reflex cannot be obtained. Left Upper Limb (UL) weakness asymmetric with respect to contralateral. Sign of extrapyramidalism with proximal stiffness of symmetrical ULs. Spontaneous position with flexion of UL over the body. No rigidity in lower limbs. No resting or positional tremor. No myoclonus. Bilateral flexor plantar skin response. No Hoffmann.

The WE was diagnosed on the basis of the clinical presentation and neuropathological analyses. The patient presented signs that met the established criteria for its detection (chronic alcohol consumption, altered mental state, oculomotor abnormalities and cerebellar dysfunction). Then, neuropathological analyses performed by responsible personnel of the Pathological Anatomy Unit in the hospital, confirmed the diagnosis of WE.

**Negative control case:** 53-year-old woman. Personal history: ex-smoker; bilateral transplant for emphysema in 2007 (post-transplant complications); Hypertension; Hypercholesterolemia.

Neuropathological diagnosis: acute global ischemic anoxia; vascular sclerosis of mild intensity in intracerebral vessels.

**Positive control case: aged brain with Alzheimer's disease (AD):** 76-year-old woman. Personal history: primary progressive aphasia, logopenic subtype. Frontotemporal lobar degeneration, in follow-up. No hypertension. No DM, no DLP. No toxic habits. No known cardiopulmonary disease. Hypothyroidism.

Neuropathological diagnosis: (1) Neuropathological changes of Alzheimer's disease, according to consensus criteria in high level (A3: THAL STAGE 4; B3: BRAAK V-VI; C3: FREQUENT CERAD); (2) Amyloid angiopathy (hyaline sclerosis of small vessels); (3) Mild atheromatosis of the polygon of Willis.

#### 1.2. Immunohistochemistry (IHC) Protocol

Tissue sections of 4 μm mounted on slides were deparaffinized and rehydrated. For unmasking or antigen retrieval, sections were exposed to pressure cooker in 0.1 M sodium citrate buffer (pH = 6) for 20 minutes. After 2 washes of 5 minutes each one in commercial wash buffer, sections were blocked of endogenous peroxidases in the tissue for 30 minutes, followed by 2 washes of 5 minutes each one in commercial wash buffer again. Then, sections were incubated for one hour with the primary antibodies against the proteins of interest dissolved in antibody diluent at their appropriate dilution. Optimal dilutions of primary antibodies were adjusted and contrasted with positive/negative technical controls. Anti-TLR4 (rabbit polyclonal antibody raised against an epitope corresponding to amino acids 242-321 mapping to an internal region of TLR4 of human origin in a dilution of 1:50; TLR4 (H-80), sc-10741 Santa Cruz Biotechnology®, CA, USA), anti-MyD88 (rabbit polyclonal antibody raised against amino acids 279-296 of MyD88 of human origin in a dilution of 1:500; ab2064 Abcam®, Cambridge, UK), anti- p-NFκB p65 (27.Ser 536) (mouse monoclonal antibody raised against a short amino acid sequence containing Ser 536 phosphorylated NFκB p65 of human origin in a dilution of 1:50; p-NFκB p65 (27.Ser 536), sc-136548 Santa Cruz Biotechnology®, CA, USA) and anti-IκB-α (affinity purified rabbit polyclonal antibody raised against a peptide mapping at the C-terminus of IκB-α of human origin in a dilution of 1:50; IκB-α (C-21), sc-371 Santa Cruz Biotechnology®, CA, USA) were used.

Consecutively, sections were washed 2 times of 5 minutes each one in commercial wash buffer and incubated with Envision for mouse or rabbit during 30 minutes in humid chamber at room temperature. Again 2 washes of 5 minutes each one in commercial wash buffer were performed. Sections were developed and visualized by diaminobenzidine (DAB) and contrasted with Carazzi's hematoxylin* for 2 minutes to stain nuclei (Carazzi's hematoxylin gives a blue or dark purple color). Finally, sections were dehydrated and embedded.

#### 1.3. Images description

Neuronal cells were differentiated from glia based on morphology. Pyramidal neurons in cortex are easily identifiable by their characteristic appearance. Glial cells such as astrocytes (light colored cytoplasm and oval nucleus), microglia (smaller), and oligodendrocytes (also smaller and with a darker coloration; round nucleus of intense color).

#### 1.4. Specific composition of the Thiamin Deficient Diet administered to TDD and CA+TDD animals

| THIAMIN DEFICIENT DIET | *Teklad Custom Diet* **TD.81029 ENVIGO** |
|---|---|
| **Key Features:** Purified Diet; Thiamin; Alcohol-Extracted-Casein; Rodent | |
| **FORMULA** | **g/Kg** |
| Casein, "Vitamin-Free" Test | 191.2 |
| DL-Methionine | 3.0 |
| Sucrose | 518.6068 |
| Corn Starch | 150.0 |
| Corn Oil | 50.0 |
| Cellulose | 50.0 |
| Mineral Mix, AIN-76 (170915) | 35.0 |
| Choline Bitartrate | 2.0 |
| Riboflavin | 0.006 |
| Pyridoxine HCl | 0.007 |
| Niacin | 0.03 |
| Calcium Pantothenate | 0.016 |
| Folic Acid | 0.002 |
| Biotin | 0.0002 |
| Vitamin $B_{12}$ (0.1% in mannitol) | 0.01 |
| Vitamin A Palmitate (500,000 IU/g) | 0.008 |
| Vitamin E, DL-alpha tocopheryl acetate (500 IU/g) | 0.1 |
| Vitamin K, MSB complex | 0.0015 |
| Vitamin D3, cholecalciferol (400,000 IU/g in sucrose) | 0.0025 |
| Ethoxyquin antioxidant | 0.01 |
| This formula is a modification of AIN-76A. Vitamin-Free Test Casein (alcohol-extracted) is used to further limit vitamin contribution from the protein source. **Background thiamin levels are below the limit of detection (<0.5 ppm).** | |

| **Selected Nutrient Information**[1] | | |
|---|---|---|
| | **% by weight** | **% kcal from** |
| **Protein** | 17.5 | 18.5 |
| **Carbohydrate** | 65.8 | 69.6 |
| **Fat** | 5.0 | 11.9 |
| **Kcal/g 3.8** | | |
| [1] Values are calculated from ingredient analysis or manufacturer data | | |

#### 1.5. Assessment of liver damage:

**Nitrite ($NO_2^-$) Liver Assay.** As the stable metabolites of the free radical nitric oxide (NO·), $NO_2^-$ was measured in the liver by using the Griess method (Green et al., 1982). Liver samples were sonicated at a ratio of 1:5 (w/v) in phosphate buffer 0.05M and were centrifuged at 13000 rpm for 15 min. The supernatant was taken for assay by using Griess reagent. A solution with 1% sulphanilamide and 0.1% N-(1-Naphthyl) ethylenediamine (NEDA) mixed in equal parts (Griess reagent) was added to the supernatants and were incubated for 20 min at RT and protected from light. Standard curve was prepared with Sodium nitrite ($NaNO_2$). The nitrites were converted into a pink compound that was read photometrically at 540 nm. The nitrite levels obtained were normalized with respect to the protein concentration in the sample and expressed as µM/mg protein.

**Lipid peroxidation (MDA: malondialdehyde) Liver Assay.** MDA levels in liver tissues were determined by a modification of the method of (Das and Ratty, 1987), based on obtaining a pink chromogen by the reaction of the thiobarbituric acid (TBA) with the MDA. Briefly, liver samples were sonicated at a ratio of 1:5 (w/v) in phosphate buffer 0.05M and deproteinized with trichloroacetic acid

(12% w/v) and HCl 5M, followed by the addition of TBA (2% w/v) in NaOH 0.5N. The reaction mixture was heated at 90°C for 15 min and centrifuged at 12000 g for 10 min. The MDA-TBA adduct of the supernatant was measured spectrophotometrically (540 nm) and the MDA concentration calculated by use of a standard curve prepared with MDA tetra-butylammonium salt. The results were expressed as nmol/mg protein.

### 2 Supplementary Data

#### 2.1. Visual/observational assessment of the IHC images by *Immuno-Reactive-Score (IRS)*

**Immunoreactivity Score (IRS): the product of the Intensity and Distribution scores gives the IRS Score (semi-quantitative scale)**

| Points | Intensity score (0-3) | Distribution score (0-4) |
|---|---|---|
| 0 | - , no | (<5%) no |
| 1 | ± , weak | 5-25% |
| 2 | + , moderate | 25-50% |
| 3 | + + , strong | 50-75% |
| 4 | – | >75% |

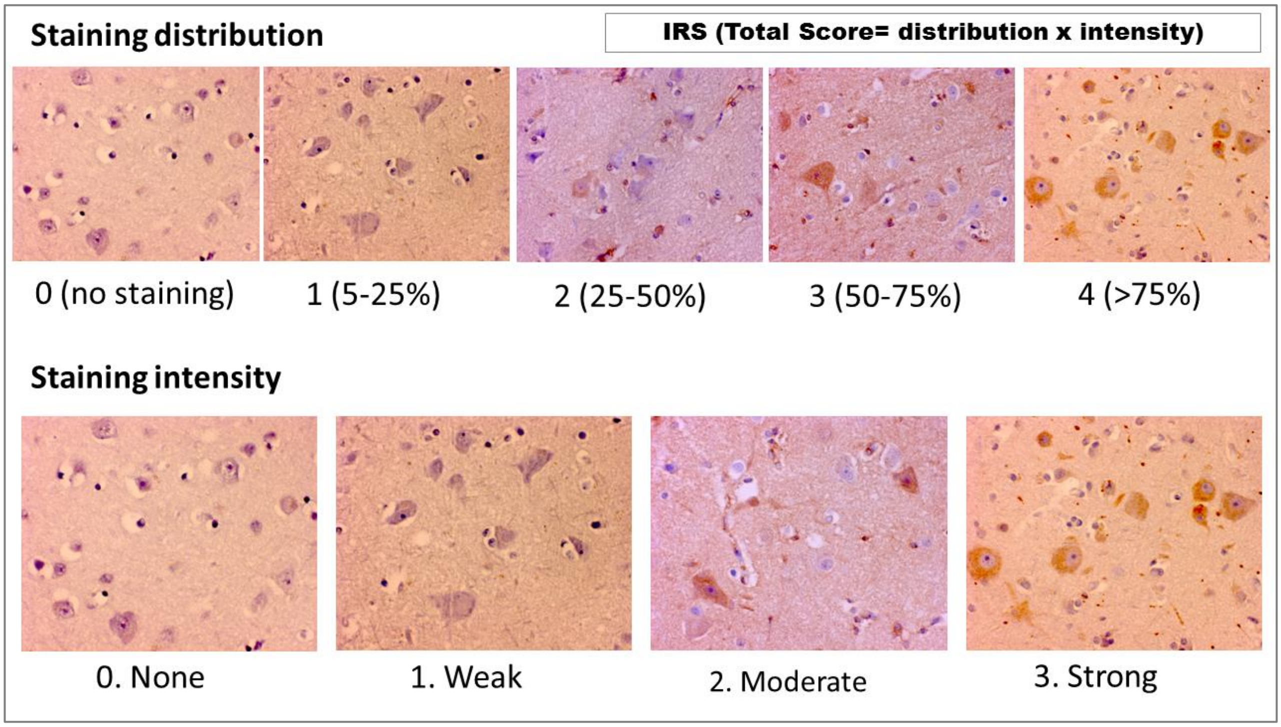

**Supplementary Figure 1.** Semiquantitative visual scoring system was used as a complementary method to automated *Fiji* to assess stained tissues, allowing the subsequent statistical analysis. The visual scoring was adapted to our study design, based on (Wang et al., 2011) and (Meyerholz and Beck, 2018). The modified IRS used here allows the observer to assign a score to the tissue changes in view of the distribution and intensity of the cellular and/or extracellular (i.e. vesicles) immunostaining. Total score was calculated from the product of the staining distribution (0-4) and intensity (0-3) sub-scores, since it increases the variation row, which gives more statistically reliable results (Fedchenko and Reifenrath, 2014).

#### 2.2. Results from manual IRS analysis of the human frontal cortex IHC images

In the cortical gray matter, significant differences between the cases were found for TLR4 staining (Supplementary Figure 2A, H=9.685, p=0.0079), finding increased TLR4 immunoreactivity in WE (p=0.0129) and AD (p=0.0045) compared with the control case.

Also within the gray matter cortex, we found higher MyD88 immunoreactivity in the AD compared to the control (p=0.0031) and WE patient (p=0.0101) (Supplementary Figure 2B, H=11.73, p=0.0028).

In the cortical white matter, also WE (p=0.0006) and AD cases (p=0.0006) showed higher TLR4 staining in this area than control (Supplementary Figure 2C, H=14.49, p=0.0007).

Interestingly, we found a pronounced elevation in MyD88 immunoreactivity in the cortical white matter of WE patient compared to control (p=0.0002) and AD cases (p=0.0002) (Supplementary Figure 2D, H=17.26, p=0.0002).

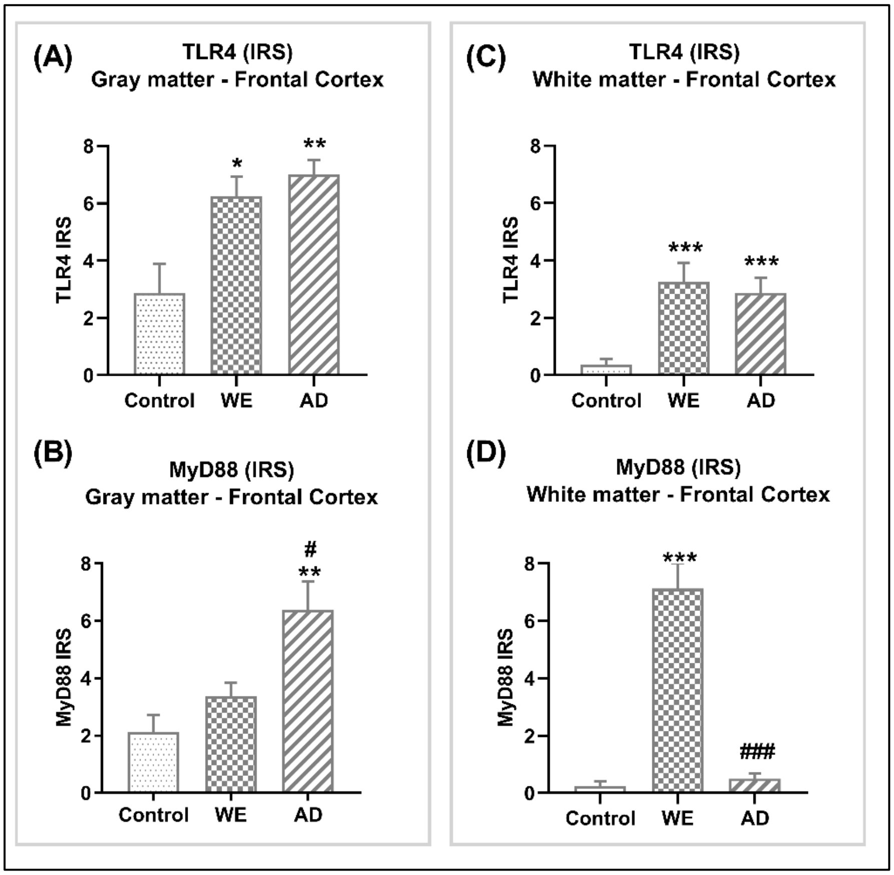

**Supplementary Figure 2. Results obtained by manual IRS processing of IHC images for TLR4 and MyD88 antibodies in the frontal cortex of control, WE and AD cases.** For this analysis, same eight separate images taken from the same tissue slide used for *Fiji* processing were employed, and final score was obtained by averaging these values for each patient. Images were assessed by a blind observer to the clinical condition (*WE: alcohol-related Wernicke's encephalopathy; AD: Alzheimer's disease*). The differences between the cases were analyzed using the non-parametric Kruskal-Wallis test followed of paired comparisons by Mann Whitney test. Mean ± S.E.M. Different from control: *p < 0.05, **p < 0.01, ***p < 0.001; different from WE: #p < 0.05, ###p < 0.001.

**Correlations between manual and automated measurements**

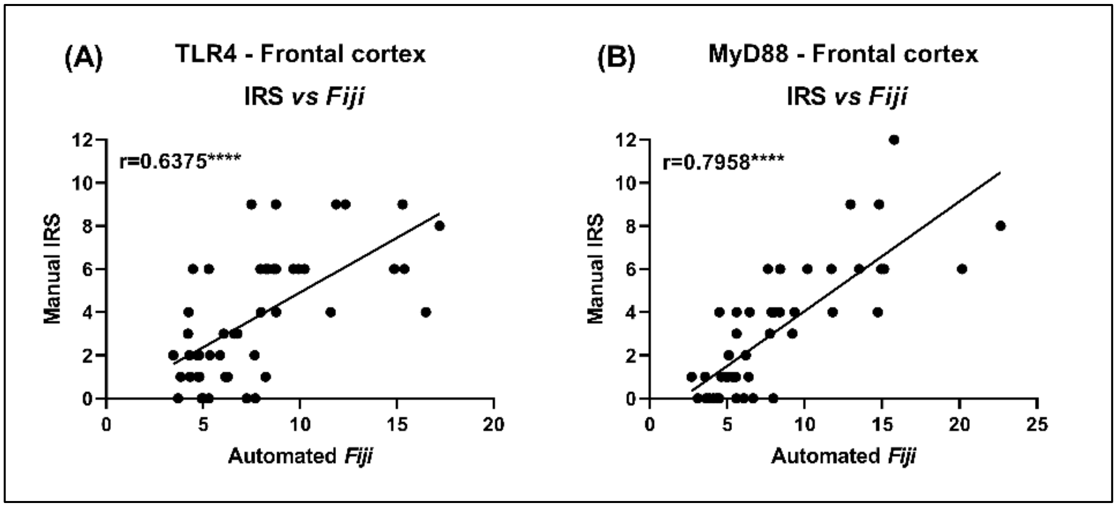

**Supplementary Figure 3.** Comparison of manual and automated scores. (A, B) Manual scores are on the y-axis, automated scores are on the x-axis. Correlation and linear regression analyses of overall scores for (A) TLR4 and (B) MyD88 of the frontal cortex. *Pearson's coefficient correlation=r;* Significant correlations (****p<0.0001).

| ***IRS vs Fiji* - Frontal cortex** | | | | |
|---|---|---|---|---|
| *Marker* | *r* | *Slope* | *Constant* | *p value* |
| **TLR4** | 0.6375 | 0.5073 | -0.1625 | <0.0001 |
| **MyD88** | 0.7958 | 0.5087 | -1.036 | <0.0001 |

**Supplementary Table 1.** Correlational and linear regression analyses between IRS and *Fiji* methods in the frontal cortex measures. *[Pearson's coefficient correlation r; Slope and constant from the linear regression analyses, p-value for both correlation and linear regression].*

**2.3. Results of other markers studied in the frontal cortex and cerebellar hemisphere of animals by Western blot analysis**

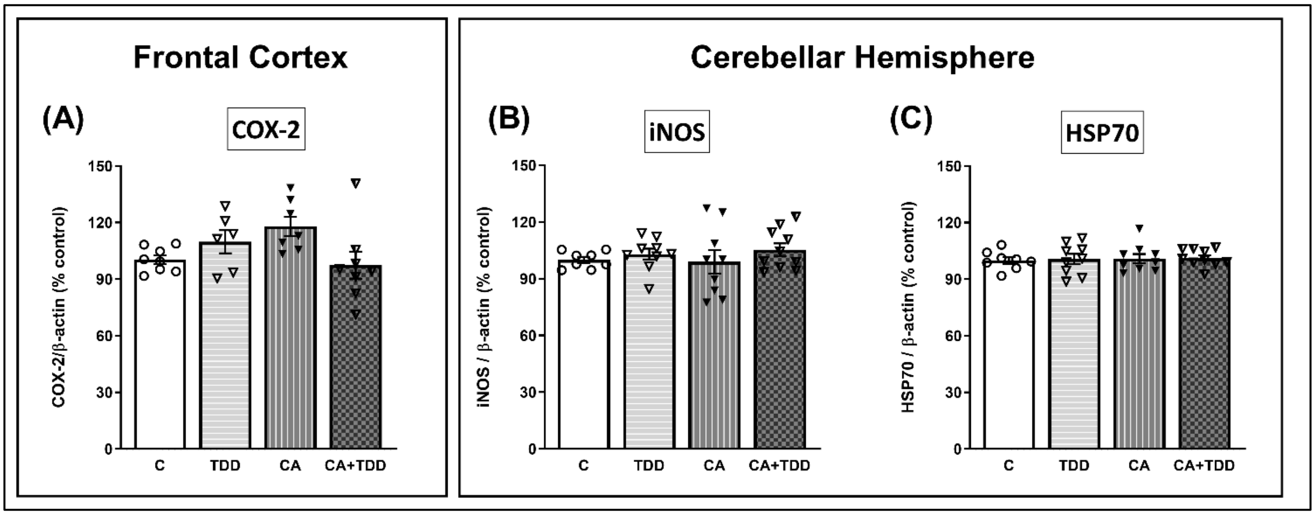

**Supplementary Figure 4. Expression of COX-2 enzyme in the frontal cortex, and iNOS enzyme and HSP70 in cerebellar hemisphere of TDD, CA and CA+TDD-treated rats.** Graphs indicate protein levels of (A) COX-2, (B) iNOS, (C) HSP70 markers by Western blot; data of the respective protein of interest were normalized by β-actin and expressed as a percentage of change versus the control group. Mean ± SEM (n = 8-10). Two-way ANOVA. Since the combined CA+TDD treatment mimics better the human case of alcohol-induced WE, this group was also compared with the C group by Unpaired Student's t-test or Mann-Whitney (CA+TDD *vs* C).

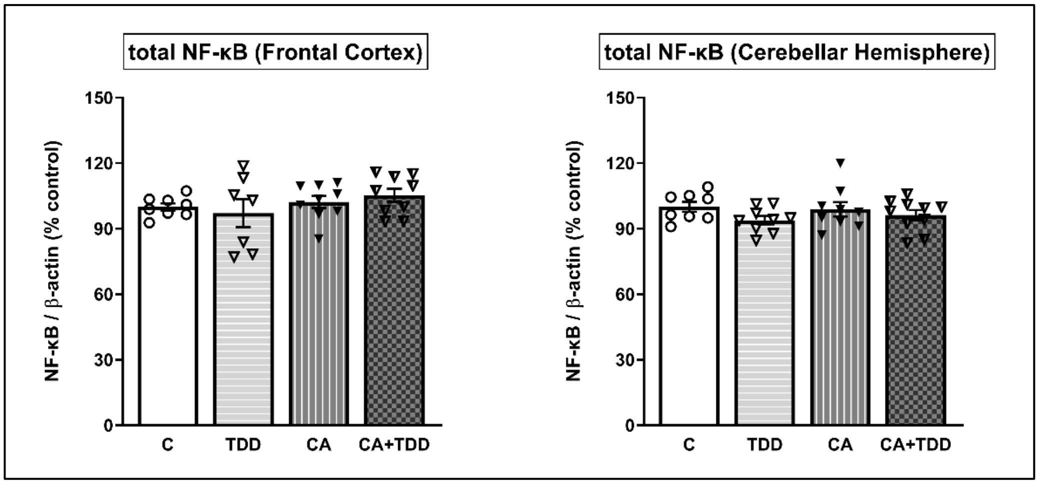

**Supplementary Figure 5. Expression of the total NF-κB factor in the frontal cortex and in cerebellar hemisphere of TDD, CA and CA+TDD-treated rats.** Graphs indicate protein levels by Western blot; data of the respective protein of interest were normalized by β-actin and expressed as a percentage of change versus the control group. Mean ± SEM (n = 8-10). Two-way ANOVA. Since the combined CA+TDD treatment mimics better the human case of alcohol-induced WE, this group was also compared with the C group by Unpaired Student's t-test or Mann-Whitney (CA+TDD *vs* C).

##### 2.4. Results of liver status check in the animals by measuring the hepatic nitrites and malondialdehyde (MDA) levels.

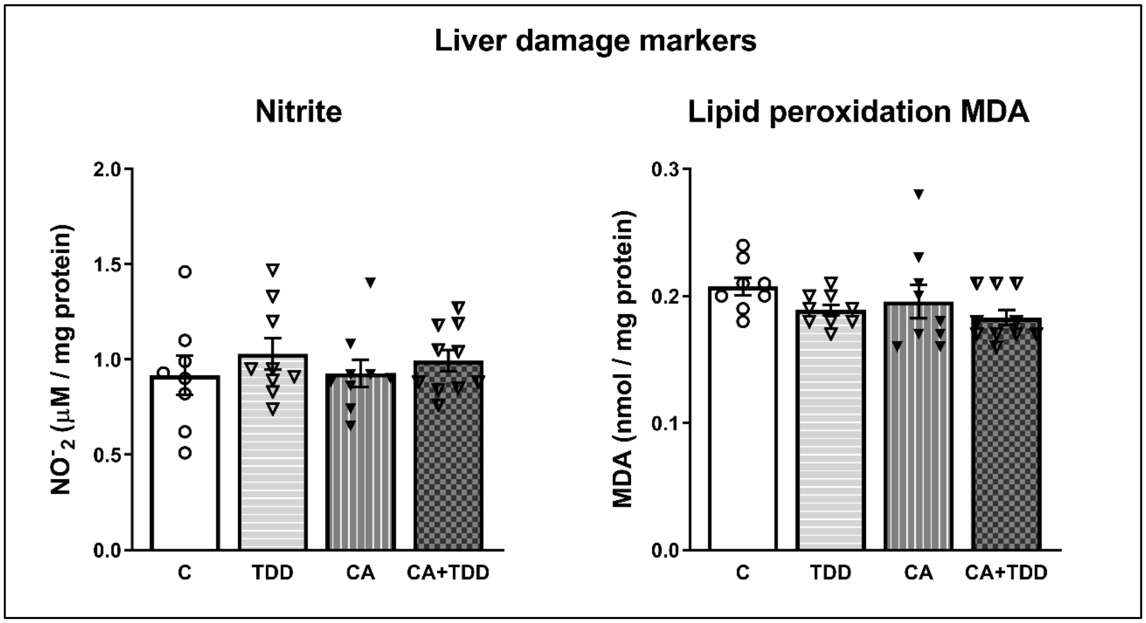

**Supplementary Figure 6. Nitrites ($NO_2^-$) and MDA measured in the liver of the rats with no signs of hepatic damage detected.** Mean ± SEM (n = 8-10). Two-way ANOVA. p>0.05, n.s.

The levels of hepatic MDA and $NO_2^-$ showed no significant changes by any of the treatments (Supplementary Figure 6, p>0.05, n.s.). Thus, the results suggest that the protocol of chronic alcohol consumption used in this study did not produce oxidative damage in the liver in the long-term, since both the nitrite and the MDA levels, indicative of nitrosative stress and lipid peroxidation, respectively, showed no significant changes in the CA and CA+TDD animals versus controls.
